# Functional coherence of bacterial ecospecies

**DOI:** 10.1101/2025.06.27.661923

**Authors:** Sarah L. Svensson, Huiqi Wen, Zhizhou Jia, Zhe Xie, Wenxuan Xu, Hui Fu, Chao Yang, Yanjie Chao, Daniel Falush

## Abstract

Understanding the molecular and evolutionary basis of adaptation is a central challenge in biology. We have recently identified bacterial “ecospecies”: strains that share a well-mixed gene pool with the rest of the species in most of the genome, but carry nearly fixed differences in ∼100 coadapted genes. We hypothesized this structure reflects ecological selection, but ecospecies have so far been characterized only genomically. Here, using “Molassodon” of the marine bacterium *Vibrio parahaemolyticus,* we provide first experimental evidence linking a multilocus ecospecies genotype to a laboratory phenotype. We find that Molassodon swimming in viscous environments is enhanced by differentiated lateral flagella that no longer drive surface swarming. Using Tn-seq, RNA-seq, and molecular genetics, we link differentiated Type VI secretion and nutrient uptake genes to the swimming phenotype, potentially invoking a natural adaptation related to “hunting”. We also identify a convergent genotype also linking lateral flagella and nutrient uptake, suggesting ecological differentiation in *V. parahaemolyticus* occurs along recurring phenotypic axes and involves borrowing traits from a broader *Vibrio* gene pool. Together, our results provide first experimental evidence that ecospecies are functionally coherent units and illustrate how they can provide a window into the complex adaptations that shape bacterial ecology.

## Introduction

Complex adaptations such as the flight of birds, the bacterial flagellum or the feeding strategies of orcas ^1^ are amongst the most reliable sources of human wonder. However, establishing the genetic basis of these adaptations and the steps involved in their evolution is typically challenging due to the large number of genes and long time periods involved. In most cases intermediate genotypes are not available, limiting our ability to investigate the genotype to phenotype map in any detail. One focus for investigation has been evolution that happens repeatedly, such as marine sticklebacks colonising rivers. Within each river, similar combinations of alleles conferring freshwater-adaptive traits are assembled onto otherwise unrelated genetic backgrounds, making it possible to identify regions of the genome responsible for adaptation ^2^.

To gain general insights into the process of complex adaptation, bacterial ecospecies are a promising new subject for investigation. A bacterial species with two ecospecies has a single-well mixed gene pool in most of the genome, but nearly-fixed differences between ecospecies at a small number of genes ^3^. We have proposed that the differences between ecospecies is generated and maintained by natural selection. High rates of homologous recombination mean that the ecospecies genotype is found on diverse genetic backgrounds. Consequently, analysis tools such as fixation indices (Fst) used for sticklebacks and other eukaryotic ecological genomics model organisms can be applied to identify differentiated regions, whose adaptive consequences can be further investigated ^2^.

For example, *Helicobacter pylori* colonizes the stomachs of humans and occasionally other mammals. The Hardy ecospecies is differentiated from typical Ubiquitous *H. pylori* at 100 of approximately 1500 genes and a handful of accessory genes ^3^. Based on the diets of humans and animals from which Hardy is isolated from as well as functional annotation of the differentiated genes, we have proposed that Hardy is a complex adaptation to host carnivory, a hypothesis that requires further functional validation. Analysis of the pattern of differentiation is consistent with Hardy evolving progressively within the stomachs of ancestors of modern humans and then dispersing around the world with us. Rampant recombination during mixed infections mean that traces of clonal descent within *H. pylori* genomes disappear within a few hundred years ^4^, but characteristic differences between Hardy and Ubiquitous strains have been maintained for at least three hundred thousand years and across diverse *H. pylori* ancestries ^3^, implying stable strong selection to maintain ecospecies differentiation at each site, in several human populations.

Some strains of *Vibrio parahaemolyticus* can also briefly colonize humans, leading to gastroenteritis ^5^. However, unlike *H. pylori, V. parahaemolyticus* is mostly free-living in oceans worldwide but is also associated with diverse marine animals, together with other *Vibrio* species. This broader environmental niche allows a wider source for its gene pool than *H. pylori*, which rarely comes into contact with other *Helicobacter* species. Our own broader population genomic analyses of *V. parahaemolyticus* have revealed extensive recombination as well as differentiated oceanic gene pools that have recently been disrupted by human activity ^6–8^.

Exploiting the high recombination rate and large effective population size in *V. parahaemolyticus*, we used genome-wide epistasis scans (GWES) to detect signals of non-randomly associated, coadapted loci, which we termed eco-LD: linkage disequilibrium driven by ecological selection rather than physical linkage or geographical population structure ^9^. Hierarchical clustering based on these interactions identified EG1a, a distinct, globally distributed group differentiated from other *V. parahaemolyticus* at ∼100 genes, and most strikingly those encoding lateral flagella. However, preliminary phenotyping analysis with a handful of strains did not reveal significant differences between different ecogroups ^10^.

Here, we demonstrate that the EG1a multilocus genotype represents a genetically and functionally coherent ecospecies of *V. parahaemolyticus,* which we now term Molassodon based on coadaptation of genes related to motility and hunting in viscous environments (molasses + Greek “tooth”), together with experimental results described below. Molassodon is differentiated from “Typical” *V. parahaemolyticus* by only 48 core and 73 accessory genes, many of which may have been acquired via introgression. We show that Molassodon is old, stable and genetically distinct from clonal, geographic, or phylogenetically defined ecotype structures.

We also identify a distinct, coherent laboratory phenotype for Molassodon: unlike Typical *V. parahaemolyticus*, it induces its differentiated lateral flagella in viscous media, where it swims faster. On the other hand, it does not swarm on surfaces. RNA-seq and Tn-seq (transposon sequencing) established the genetic basis of the positive swimming phenotype and linked it to further differentiated traits, including genes involved in nutrient uptake and Type VI secretion. This linkage, combined with further phenotyping of Molassodon strains, suggests that the adaptive advantage encoded by Molassodon loci could be related to active foraging (“hunting”) of bacterial prey in viscous environments. Finally, we identify a convergent, intermediate genotype that provides additional support for the functional link between motility and nutrient acquisition in *V. parahaemolyticus* ecology. These genomic and experimental findings reinforce the ecospecies concept as a meaningful organizing principle, and open new avenues for using ecospecies to explore the genetic basis of complex adaptations and the ecological factors driving them in bacteria.

## Results

### The multilocus genotype identified previously by epistasis analysis is an ecospecies, Molassodon

We previously identified a multilocus genotype in *Vibrio parahaemolyticus* (EG1a) using a genome-wide epistasis scan (GWES) for non randomly-associated, coadapted loci ^10^. This involved pairwise Fisher’s exact tests between variants in a non-redundant genome dataset from the VppAsia population. This identified a large network of coadapted genes, and hierarchical clustering revealed several strain clusters, of which EG1a was the most distinct. Since EG1a is differentiated in many genes but also present in several geographic populations/genetic backgrounds, we hypothesized that it is a bacterial ecospecies, comparable in structure to the *H. pylori* Hardy ecospecies ^3^.

We performed the same GWAS analysis that we performed to characterize Hardy^3^. In addition, since deeper functional annotation of Hardy genes allowed us to make a compelling hypothesis about the phenotypic/ecological difference of the ecospecies, we carefully inspected the putative function of differentiated genes. Based on this additional genomics analysis described below, we now designate EG1a as an ecospecies, which we refer to as “Molassodon” throughout the manuscript for clarity [based on functional annotation and experiments also described below that suggest the ecospecies might be adapted to hunting (-don, tooth) in viscous (molasses) environments]. We refer to all other strains as “Typical”.

The ecospecies genetic signature we observed for *H. pylori* Hardy was small islands of near-fixed differentiation within a freely mixing genome ^3^. The signature was detected in several geographic populations, and differentiated genes showed a distinct evolutionary trajectory from the rest of the genome. To confirm that EG1a is an ecospecies, we performed a similar core-genome GWAS (genome-wide association study), comparing Molassodon and Typical strains using a larger non-redundant genome dataset spanning all four populations (VppAsia, VppX, VppUS1, VppUS2) ^6,9^. Strains were independently classified as Molassodon/Typical by GWES essentially as performed previously on a non-redundant dataset of 1550 strains from four geographic populations (**Fig. S1A, B & S2A; Methods**) ^10^. We did not observe any population structure between ecospecies using FineSTRUCTURE ^11^ (**Fig. S2B**).

Only ∼15 core genome regions showed near-fixed differences between Molassodon and Typical *V. parahaemolyticus* (*F_ST_* > 0.9; **Fig. 1A**), corresponding to 46 differentiated core genes (≥5 significant SNPs; −log_10_ > 10; **Tables S1–S4**). These regions were distributed across the two chromosomes of Molassodon and Typical strains (**Fig. S3**). Differentiation decayed to near zero within a few hundred base pairs of these loci (**Fig. 1B**; **Fig. S4**), as observed previously for *H. pylori* Hardy ^3^. Thus, Molassodon differs from Typical strains at only a small number of loci, with little differentiation elsewhere in the genome, consistent with ecospecies structure.

**Figure 1.**
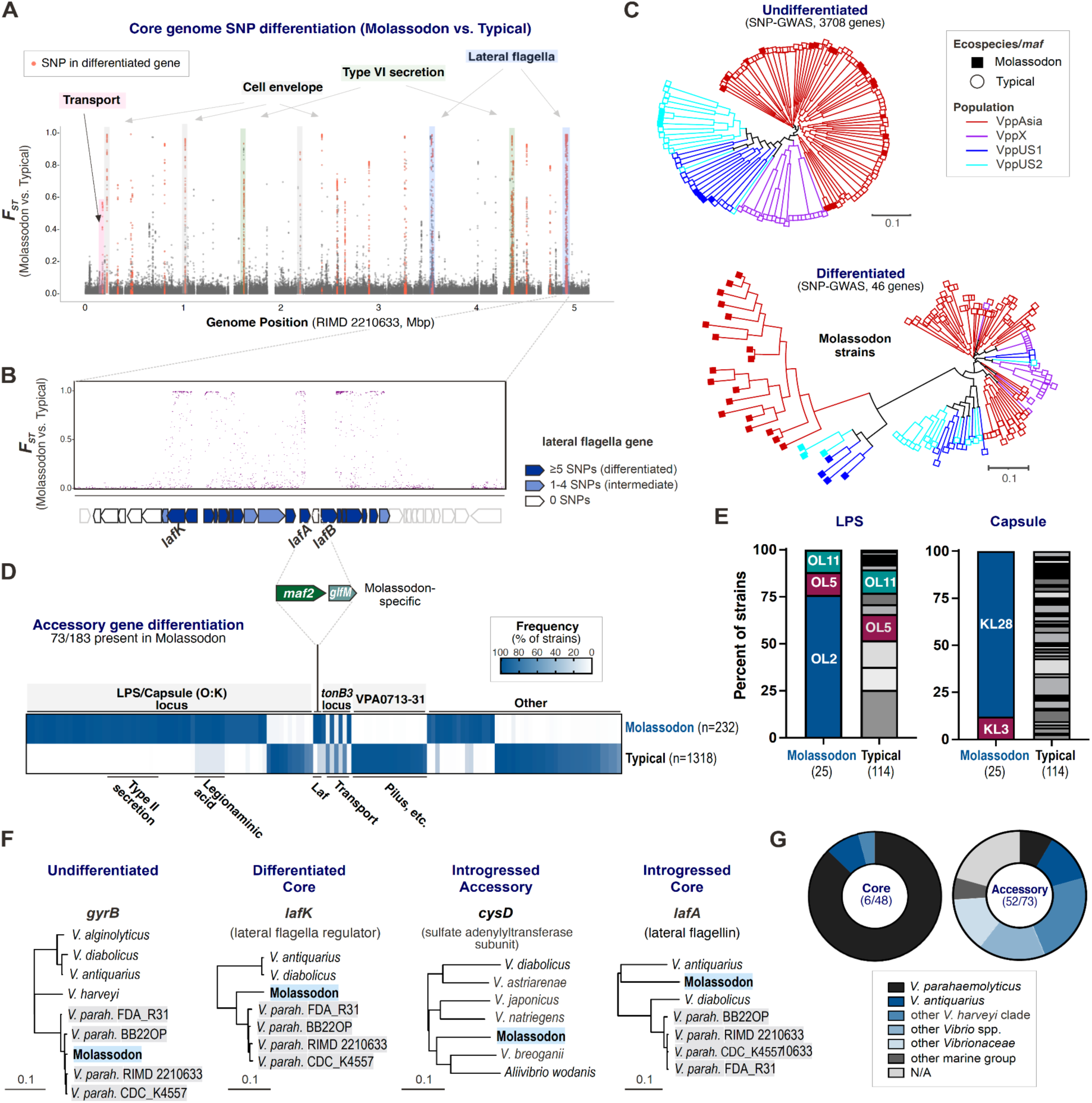
The Molassodon ecospecies genotype. **(A)** Manhattan plot for core genome SNP (single-nucleotide polymorphism) GWAS of Molassodon (n=223) vs. Typical (n=1318) strains. Significant SNPs: *F_st_* (fixation index) ≥0.5 and -log10(p) ≥10. Only those in differentiated genes (5 or more SNPs) are labeled in red. Key functions relevant to this study are highlighted. **(B)** Zoom-in on indicated lateral flagella region in *F_sy_* plot in panel A**. (C)** Approximate Maximum Likelihood trees for Undifferentiated (*top*, 3708 genes) and Differentiated genes (*bottom,* 46 genes) identified in core genome SNP GWAS. A subset of strains with representatives of each group of interest (population, ecospecies) was used. Differentiated genes: at least five differentiated SNPs. Undifferentiated genes: no SNPs with *F_st_* ≥0.05. Note that two highly differentiated core genes identified in the pan-GWAS are not included. Branch colours: geographic population. Tip shape: ecospecies (Typical/Molassodon) genotype. **(D)** Frequency of accessory genes identified as significantly differentially distributed between Molassodon and Typical strains via pangenome GWAS. Significant genes: *F_ST_* ≥ 0.5 and -log_10_(p) ≥ 10, also requiring >90% or <10% frequency in Molassodon. Genes are ordered by putative function. **(E)** Distribution of predicted serotypes for the reduced set of 136 genomes (Fig. 1C). Serotypes present in Molassodon strains are coloured. Serotypes were predicted from genome sequences with Kaptive ^12,13^. **(F)** Representative trees for KEGG matches for undifferentiated (*gyrB*) and representative example genes for the three types of Molassodon-differentiated genes: 1) Core genes with closest match in other *V. parahaemolyticus* strains, but with an extended branch length; 2) Accessory genes with signs of introgression (*i.e.*, closest match outside of *V. parahaemolyticus*); 3) Core genes with signs of introgression. Grey: Typical *V. parahaemolyticus*. Blue: Molassodon. **(G)** Taxon of best KEGG ortholog (by % amino acid identity) for Molassodon-present accessory and core genes. N/A: no match above 40%.

Phylogenies based on *H. pylori* ecospecies-differentiated core genes were markedly distinct from those based on only undifferentiated genes: the former clustered Hardy strains together on a single long branch, whereas the latter tree showed them dispersed among Ubiquitous strains according to their geographic population ^3^. We found that this pattern is recapitulated for *V. parahaemolyticus* Molassodon. A phylogeny based on undifferentiated genes (no SNPs with *F_ST_* ≥ 0.05; n = 3,708; **Table S3**) was star-like (**Fig. 1C**, *top*), as expected for highly recombining *V. parahaemolyticus*, and largely reflected geographic population structure ^6^. Molassodon strains, which were detected in three of four geographic populations, were distributed among Typical strains, and according to their assigned geographic population. This indicates that Molassodon is largely undifferentiated from Typical strains outside of coadapted loci. In contrast, a phylogeny based on the 46 ecospecies-differentiated core genes clustered all Molassodon strains together on a single long branch, independent of their population assignment (**Fig. 1C**, *bottom*). Thus, as we observed for *H. pylori*, the small handful of differentiated ecospecies genes have a distinct evolutionary trajectory from the rest of the genome, despite extensive within-species gene flow. Based on this, we concluded that the multilocus EG1a genotype identified by GWES represents a distinct ecospecies, which we now term Molassodon based on functional characterization.

### The Molassodon ecospecies is much older than identifiable clonal lineages

In addition to the pandemic clonal lineage that causes the majority of human infections, our global *V. parahaemolyticus* dataset has sets of strains that are more distantly related, but still show signs of common descent. For example, we detected two “deep clones” of 18 and 21 strains visible in the phylogenetic tree of non-redundant strains used for GWAS, all assigned to the VppAsia population (**Fig. S1A**). The distribution of genetic distances between strains in each clone (median 41,936 and 49,512, respectively, **Fig. S5A, B**) is clearly lower than between unrelated strains in the VppAsia population (median 54,875) and is consistent with them sharing an average of approximately 10% of their genome by direct descent. Using estimates of the recombination rate from contemporary evolution of epidemic clones ^9^ and assuming that recombination falls randomly in the genome, this implies that a typical pair of strains for the 21-strain clone shared a common ancestor around 6-7,000 years ago, while the 18-strain clone is younger with a common ancestor of ∼4,000 years ago (**Fig. S5B**; see **Methods** for details). The common ancestor of all 21 of the strains in the deep clone is necessarily older but making estimates for this or older events would require stronger modelling assumptions, because at this time depth recombination has erased most of the evolutionary signal.

If we apply the same approach to Molassodon, we find that the signature of shared ancestry by direct descent is tenuous or non-existent. The median SNP distance between VppAsia Molassodon strains is 54,440 SNPs (**Fig. S5B**), which is only 435 SNPs fewer than the median SNP distance between VppAsia strains as a whole. If this difference is used to estimate shared ancestry, it implies that around 1% of the genome of pairs of Molassodon strains is shared by direct descent, with the genome being overwritten around 5-10 times on average by recombination. This would be consistent with the average pair of strains sharing ancestors 14,000 years ago (using the median) or 26,000 years ago (using the mean). However, the assumption of the model that recombination is random clearly breaks down, because we have seen above that natural selection has preserved differences between Molassodon and Typical strains in ∼40 genes out of 4100 in the core genome. Thus, the data is also consistent with substantially older pairwise coalescence times. Furthermore, there are 220 non-redundant Molassodon strains in VppAsia, meaning they are ∼10-fold more numerous in the dataset than the deep clones, as well as Molassodon strains in two other geographic populations (VppUS1 and VppUS2). Since all of these lineages need to coalesce to get to the common ancestor of Molassodon, this happened at an even greater time depth, but once again, this date could only be estimated by making strong modelling assumptions, as SNP data provides no information on these older events.

Also in line with an older and distinct structure from clonal lineages or acquisition of a single block of accessory genes as reported for, e.g., virulence-associated evolution, we observed many genome rearrangements in Molassodon compared with Typical strains, especially in Chromosome 1 (**Fig. S3**). This also showed that the location of differentiated core genes was distributed across both chromosomes in both Typical and Molassodon.

The pattern of core-genome differentiation for the Molassodon ecospecies is also distinct qualitatively from other population structures such as the detected clones or geographic populations. Consistent with the homogenizing effect of recombination over long time periods, Molassodon is undifferentiated from Typical strains for more than 90% of the genome and shows strong differentiation in less than 1% of the genome (**Fig. 1A**). Differentiation plots for the deep clones or Molassodon strains from the same geographic population vs. Typical are strikingly different, with differentiation spread across the genome (**Fig. S6**). Likewise, geographic population differentiation is genome-wide (**Fig. S7**), rather than focused in small islands representing ∼1% of the genome. Therefore, Molassodon is distinct from *V. parahaemolyticus* clones and geographic populations.

### Accessory and introgressed traits also distinguish Molassodon and Typical strains

The *H. pylori* Hardy ecospecies genotype includes both core-gene alleles as well as accessory genes (*e.g*., an iron-cofactored urease) ^3^. We therefore expanded on the 46 differentiated core genes by performing a GWAS for accessory gene presence/absence, identifying 73 genes enriched and 65 depleted in Molassodon strains vs. Typical (**Fig. 1D**; **Table S4**). Two of the identified accessory genes are actually core genes in the *tonB3* locus (VP0162/0163) that are sufficiently divergent to be called as “accessory” genes (**Fig. S8, S9**); thus, we added these to our core genome list to make a final count of 48.

As noted previously ^10^, most Molassodon-associated accessory genes (59/73; **Table S5**) are concentrated in a large genomic block. We now note that this block corresponds to the LPS/capsule locus, which underlies O:K serotyping in *Vibrio* spp. ^14^ (**Fig. S10**; **Table S6**). Despite generally high serotype diversity in the analyzed dataset for Typical strains, Molassodon strains carry very few O or K types (OL2/5/11; KL3/28) (**Fig. 1E**). Previous work has shown that KL3/28 types are highly divergent and may be the result of two insertion events from other species ^15^. Our inspection likewise suggests that several genes in this locus derive from several introgression events, with closest matches predominantly in other Vibrionaceae, most frequently *V. antiquarius* (**Table S5**).

We identified three evolutionary patterns among Molassodon-differentiated genes (**Fig. 1F, G; Fig. S8**). First, most core genes evolved via point mutation, with the Molassodon version branching together with other *V. parahaemolyticus* strains, albeit often with a slightly extended branch length (*e.g*., *lafK*). Second, most Molassodon accessory genes (52/73) bear signatures of introgression from other species, as they have closest homologs outside of *V. parahaemolyticus* (*e.g.*, *cysD*). Finally, we also identified six core genes with evidence of allelic exchange with other species (*V. antiquarius* and *V. rotiferianus*), including *lafA* and genes in the *tonB3* locus (**Figs. 1F**, **S8B**; **Table S3**). Notably, core gene introgression has been reported previously in *Vibrio* ^16^. In sum, this implies that a large proportion of the multigene Molassodon genotype has been acquired from outside of *V. parahaemolyticus*.

Overall, the above systematic analyses 1) confirm that the multilocus genotype EG1a is an ecospecies, which is distinct from a population or a clone, 2) that Molassodon is differentiated in only 48/∼4800 nearly fixed core genes and over 70 accessory genes, and 3) that a significant portion of the genotype may have come from outside of *V. parahaemolyticus.* Having characterized the genotype, we moved on to understanding the Molassodon phenotype and the connection of coadapted genes with it.

### Molassodon strains do not swarm despite inducing lateral flagella

Our strongest clue to a tractable phenotypic difference between Molassodon and Typical ecospecies was motility, based on the strong differentiation of lateral flagella genes (**Fig. 1A**). In *V. parahaemolyticus*, lateral flagella are environmentally regulated and enable swarming motility on surfaces, in contrast to the constitutively expressed polar flagellum that mediates swimming in liquids. Closer examination revealed that both structural components and multiple regulators of the lateral flagellar system are differentiated in Molassodon strains (**Fig. 2A**). However, the most differentiated core gene based on the number of SNP differences encodes LafK, a key regulator of lateral flagella. This suggested there might be not only a functional divergence in Molassodon lateral flagella, but also a difference in how they are regulated in response to the environment. We previously compared motility of a small number of EG1a and non-EG1a (Molassodon and Typical) strains, which did not reveal a significant difference ^10^.

**Figure 2.**
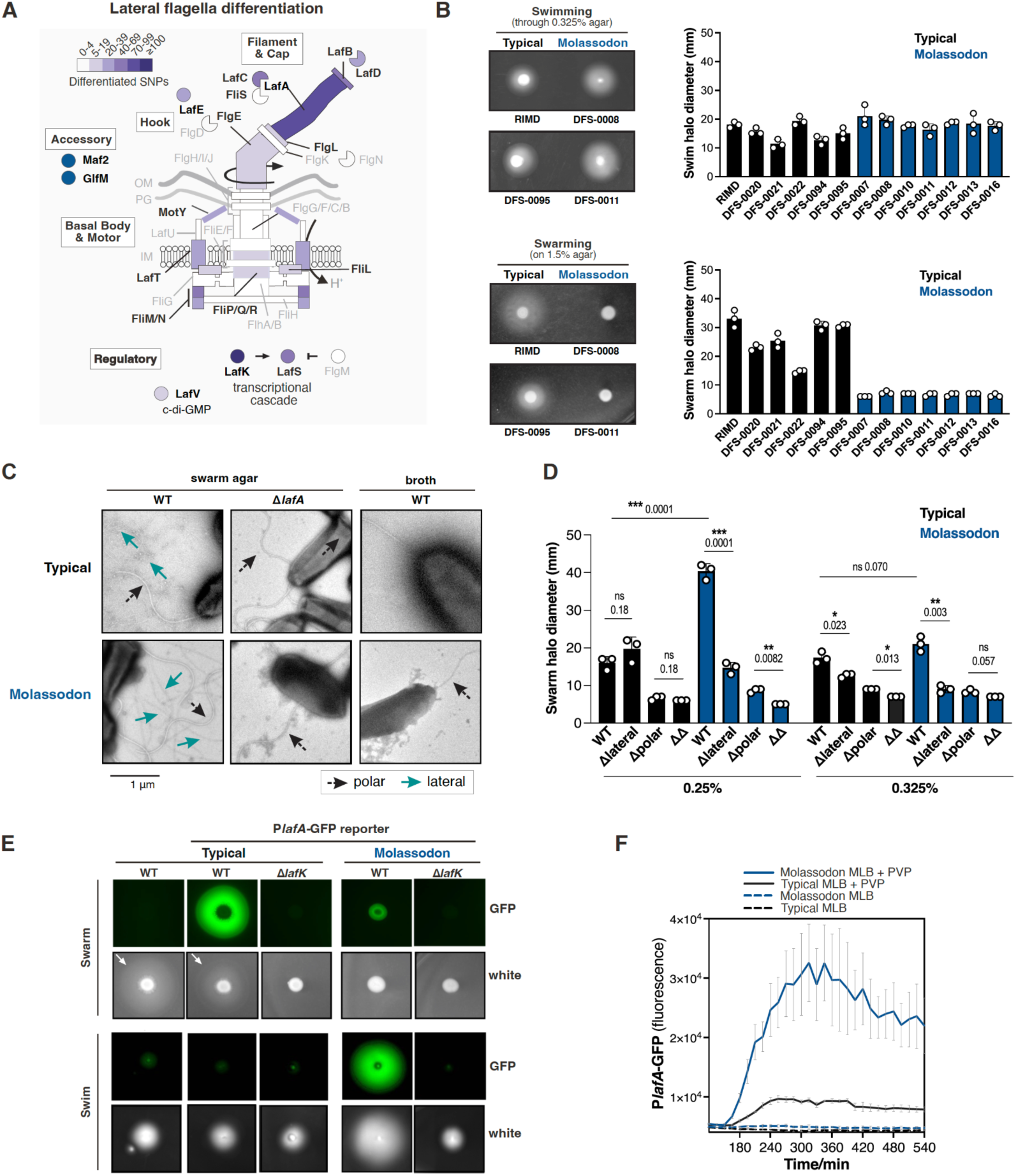
Molassodon uses lateral flagella for swimming in viscous media rather than swarming. **(A)** Schematic of lateral flagella differentiation in Molassodon strains. Differentiated: *F_st_* ≥0.5 and -log10(p) ≥10; Molassodon vs Typical GWAS. **(B)** *Left*: Representative swim/swarm halos for Typical and Molassodon strains. *Right*: Halo measurements. Representative of two independent experiments. **(C)** Transmission electron micrographs of Typical and Molassodon strains on swarm agar. Δ*lafA*: deletion of lateral flagellin gene (VPA1548). Typical WT was harvested from the outside of swarm halos; Molassodon WT and Δ*lafA* for both ecospecies were recovered from the macrocolony that forms in the absence of swarming (*e.g.*, Molassodon, panel B). Arrows: polar (black) or lateral (green) flagella. **(D)** Swim assay in low concentration (0.25%) agar for Typical and Molassodon WT or polar/lateral flagella deletion strains. Δlateral: Δ*lafA*, lateral flagellin. Δpolar: Δ*fliS*, polar flagellin chaperone. Neither: Δ*lafA* Δ*fliS*. *** *p* < 0.001, **** *p* < 0.0001, ns - *p* > 0.05, Student’s *t*-test. **(E)** Activity of *lafA* promoter reporters (∼200 nt upstream of the transcriptional start site) in Molassodon/Typical backgrounds on swarm agar (*top*) and in swim agar (*bottom*). Representative of at least three independent experiments. **(F)** Expression of P*lafA* reporters in Molassodon and Typical in the absence (MLB) and presence (MLB+3% PVP, polyvinylpyrrolidone). Error bars: standard deviation for three strains of the same ecospecies.

Laboratory motility assays using protocols more widely used for *V. parahaemolyticus* ^17^ revealed a striking phenotypic divergence between Typical and Molassodon strains. We compared swimming (through soft agar, driven by polar flagella) and swarming (migration on a solid agar surface driven by lateral flagella) in natural Molassodon (n=7) and Typical (n=6) isolates. Molassodon strains formed modestly larger motility halos than Typical strains in 0.325% swim agar (**Fig. 2B**; **Fig. S11A**), but strikingly, showed no surface migration on swarm plates and instead grew as apparently immobile macrocolonies. Despite the absence of swarming in Molassodon, RNA-seq confirmed surface induction of lateral flagella genes, albeit at ∼10-fold lower levels than in Typical strains (**Fig. S11B**; **Table S8**). Molassodon strains could also still assemble lateral flagella when grown on surfaces despite lacking macroscopic movement (**Fig. 2C**).

### Molassodon uses lateral flagella for swimming through viscous conditions

We hypothesized that Molassodon strains might use their differentiated lateral flagella for an alternative function. To first confirm that Molassodon lateral flagella could in fact drive movement of any sort, we constructed mutants expressing only polar or only lateral flagella and measured migration in more permissive 0.25% soft agar (vs. the previous concentration of 0.325%). Both Molassodon and Typical strains expressing only lateral flagella (Δ*fliS*; VP2254, polar flagellin chaperone) could still form small swim halos, which were abolished by additional deletion of the major lateral flagellin gene *lafA* (**Fig. 2D**). These results demonstrate that Molassodon lateral flagella should be functional to drive motility.

This experiment unexpectedly revealed a “positive” Molassodon motility phenotype (*i.e*., stronger than in Typical and not a “defect”) that was driven by lateral flagella. First, we noticed that Molassodon strains formed 2-fold larger motility halos than Typical strains in the 0.25% soft agar (**Fig. 2D**). This was not significant in the higher 0.325% agar (**Fig. 2D**). We confirmed this enhanced swimming phenotype for the natural strains tested above (**Fig. S12A**). Strikingly, deletion of *lafA* reduced Molassodon swimming by half compared to the wild-type Typical strain, but did not affect Typical strain migration. This is consistent with the expected minimal lateral flagella expression during swimming in Typical *V. parahaemolyticus*. Moreover, it provides evidence that Molassodon might uniquely induce and use its lateral flagella in these conditions, conferring a motility advantage.

To confirm that Molassodon induces lateral flagella during swimming, we generated a transcriptional GFP reporter for the promoter of *lafA* for each ecospecies (**Fig. S12B**). As expected, Typical strains induced the *lafA* promoter only on swarm agar, whereas Molassodon activated it while swimming (**Fig. 2E**). RNA-seq of three strains per ecospecies taken from swim agar confirmed higher expression of all lateral flagella genes in Molassodon compared with Typical in this condition (**Fig. S12C**; **Table S9**), which is in line with the Δ*lafA* swimming defect in only Molassodon (**Fig. 2D**).

What could be the specific signal resulting in differential induction of lateral flagella between the ecospecies? Surface sensing and viscosity can trigger lateral flagella expression via inhibition of polar flagella rotation in *V. parahaemolyticus* ^18^, so we hypothesized that this could be one environmental trigger. To test this, we next measured the effect of polyvinylpyrrolidone (PVP), a polymer that increases viscosity, on expression of the *lafA* reporters. We found that Molassodon induced *PlafA* more strongly than Typical strains in response to increased viscosity (**Fig. 2F**; **Fig. S12D**). Together, these results reveal the first laboratory phenotype for a bacterial ecospecies associated with differentiated genes and suggest that Molassodon might uniquely induce and use lateral flagella for swimming in viscous environments.

### Functional linkage of most differentiated genes to lateral flagella-dependent swimming

Having identified, at least in the laboratory, a “positive” phenotypic difference for Molassodon that is linked to lateral flagella (swimming in viscous media), we hypothesized that we could leverage it to assay if other Molassodon-differentiated genes might also be linked to this behavior. Potentially, the other differentiated genes could be 1) hidden essential motility factors, or 2) genes that are co-expressed under the same condition because they are part of a broader, emergent phenotype representing a complex adaptation involving most of the multilocus genotype. To test these two possibilities, we used Tn-seq to identify genes required for Molassodon and Typical swimming (log_2_FC < 1 vs. shaken media) and our RNA-seq dataset from the swim condition to identify genes co-expressed with lateral flagella (log_2_FC > 1 vs. LB) in each ecospecies (**Fig. 3A; Fig. S13A**). As a control, we also measured phenotypes and expression in the Typical strain during swarming (**Fig. S13A,** *top*).

**Figure 3.**
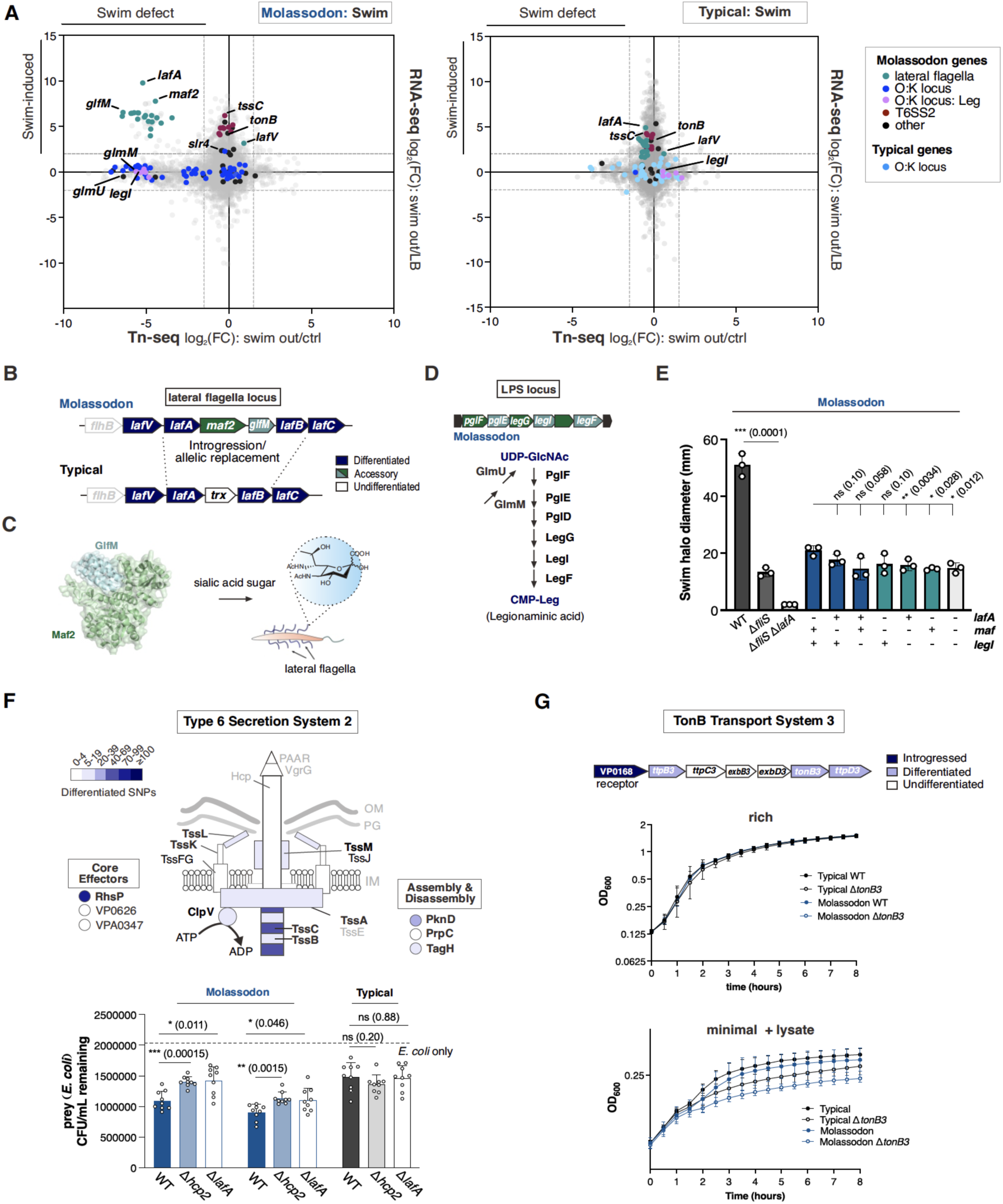
Validation and refinement of the Molassodon laboratory phenotype with Tn- and RNA-seq. **(A)** Tn-seq and RNA-seq profiles for Molassodon and Typical strains during swimming. Tn-seq: RIMD 2210633 (Typical) and DFS-0010 (Molassodon), log_2_FC Swim halo vs. Swim control. RNA-seq: mean of three Typical or Molassodon strains, log_2_FC Swim halo vs. LB. Coloured dots: Molassodon genes based on GWAS; Grey: all other genes shared by the ecospecies. **(B)** Genomic context of the Molassodon accessory lateral flagella genes *maf2-glfM* and related region in non-introgressed strains. **(C)** Predicted complex formed by Maf2-GlfM and role of the proteins in flagellin glycosylation. The complex was predicted with Alphafold Multimer at Colabfold ^22,23^ and visualized with Pymol. **(D)** Molassodon genes encoding a putative Leg (legionaminic acid) biosynthesis pathway. The pathway is based on ^24^). CMP: cytidine monophosphate. **(E)** Motility assays with *maf2* and *legI* mutants. *Left*: Epistasis analysis of the contribution of Maf (*maf2-glfM*), Leg biosynthesis (*legI*), and lateral flagella (*lafA*) on swimming motility in Molassodon. *Right*: Effect of *legI* deletion on swimming in a Typical strain. *** *p*<0.001, ** *p*<0.01, * *p*<0.05, ns: not significant, Student’s *t*-test, n=3. *fliS*: polar flagella chaperone. **(F)** *Top:* Schematic of T6SS2 with differentiated components indicated. Differentiated: *F_ST_* ≥0.5 and -log10(p) ≥10); Molassodon vs Typical GWAS. *Bottom:* Killing activity against *E. coli* BW25113 for Typical and Molassodon wild-type, *lafA*, and T6SS2 (Δ*hcp2*) mutants in M9 broth (with 2% NaCl) + 5% PVP. The colony-forming units of the *E. coli* strain were determined after 3.5 hours of incubation at 30°C and normalized to the number for the corresponding *E. coli* group without added attacker for each ecospecies. Dashed line: mean prey CFU/ml remaining without added *V. parahaemolyticus* attacker *** *p*<0.001, ** *p*<0.01, * *p*<0.05, ns not significant, Student’s *t*-test**. (G)** *Top:* Schematic of differentiated genes encoding a TonB/ExbBD transporter and receptor in Molassodon. *Bottom:* Growth curve analysis of Molassodon and Typical strains (n=3 each) and isogenic *tonB3* (VP0163) deletion strains in rich (MLB) or minimal medium (M9) with prey (*E. coli*) lysate is provided as a nutrient source.

Of the 116 Molassodon genes that we could assay with both Tn-seq and RNA-seq, 76 (66%) were either required for swimming or induced in the swim condition where only Molassodon uses lateral flagella (39 induced, 56 required, 19 overlapping; **Fig. 3A**; **Table S10**; **Fig. S13B**). In contrast, few differentiated core genes were induced or required by Typical in the swim condition (**Fig. 3A**; **Table S11**). As a control, we confirmed that Typical lateral flagella genes were required for swarming (**Fig. S13A**; **Table S12**). Overall, Molassodon genes were highly enriched in both swim-induced and swim-required sets (padj = 5.36×10⁻¹⁶, NES = 3.36; padj = 3.82×10⁻¹², NES = −2.68), indicating that this swimming phenotype, although tested in the lab, might encompass most Molassodon-differentiated genes. It follows that we may have potentially recapitulated part of the ecological scenario where Molassodon genes together provide an adaptive advantage.

To understand why the other required/co-expressed Molassodon genes might be linked to motility, we examined their functions. First, we noted that several differentiated genes in the LPS (“O”) locus, including genes for legionaminic acid (Leg) sugar biosynthesis, showed strong Tn-seq phenotypes (**Fig. S14A**). A block of K-locus genes, previously suggested to be gained via insertion event from other species ^15^, also had a strong Tn-seq phenotype. Synteny and structural similarity analysis suggested that these genes are homologous to a recently described marine bacterial S-layer and its dedicated Type II secretion system ^19^ (**Fig. S14B-D**). S-layers are proteinaceous, crystalline layers expressed on the surface of some bacteria and many archaea that can protect from predation/phages and confer unique surface properties ^20^. We confirmed via SDS-PAGE analysis that Molassodon strains express a highly abundant protein that disappears upon deletion of the putative S-layer protein gene, *slr4* (**Fig. S14E**), although we have not yet visualized an S-layer. Finally, this region also encodes a distinct ABC transporter-dependent capsule, unlike the Wzx/Wzy-dependent version of Typical strains ^15^, which also mildly affected swimming (**Fig. S14A**).

Tn-seq phenotypes for many of the O:K locus genes such as those encoding Leg biosynthesis (**Fig. 3A**) at first suggested altered cell envelope/surface characteristics might indirectly affect swimming. However, we did not see the same impact on Typical swimming (or swarming) (**Fig. 3A; Fig. S14A**). This led us to investigate a direct link between other differentiated genes and lateral flagella.

Two uncharacterized Molassodon accessory genes encoded downstream of *lafA* (now named *maf2* & *glfM*) showed a strong swimming defect in the Tn-seq experiment (**Fig. 3A; Fig. S14A**). Based on homology, predicted complex formation, and co-inheritance with the Molassodon *lafA* allele, these two genes might encode a Maf2-GlfM flagellin glycosylation system as was recently described in *Shewanella* ^21^ (**Fig. 3B & 3C**). Maf2 and GlfM likely act together to glycosylate Molassodon LafA with a sialic-acid–derived sugar. Consistent with this, the Molassodon OL2/OL5/OL11 LPS islands encode a putative pathway for the biosynthesis of the sialic acid legionaminic acid (Leg) (**Fig. 3D; Fig. S10B**), and O2 serotype LPS carries a Leg derivative ^25^. Molassodon *lafA*, *maf2*, and *legI* appear to have been imported from outside of *V. parahaemolyticus*, being closer to that found in *V. antiquarius* (**Fig. S14F**).

All six Leg pathway genes, along with two Molassodon-differentiated core genes feeding into this pathway (*glmM*, *glmU*), had Molassodon-specific Tn-seq phenotypes in the swim condition, but did not affect Typical swimming or swarming (**Fig. 3A, Figs. S13A & S14A**). Single and combinatorial mutants of *lafA*, *maf-glfM*, and *legI* phenocopied each other, producing swim halos similar to Δ*lafA*, which provides support that these genes act in a shared pathway affecting Molassodon lateral-flagella-dependent swimming (**Fig. 3E**). Therefore, a functional connection to lateral flagella via a role in generation of a sialic acid substrate for Maf explains Tn-seq swimming phenotypes for several additional genes encoded outside the lateral flagella loci, and especially at the LPS locus. Because Maf is presumably not functional without genes encoding the biosynthesis pathway of its substrate, this also suggests that Leg biosynthesis, and a dedicated role in something like LPS modification, preceded the acquisition of the Maf glycosylation function during Molassodon evolution. This provides a plausible model for how at least part of the multilocus ecospecies genotype may have been assembled in a step-wise fashion.

### Type VI-mediated killing and nutrient uptake are also functionally linked to Molassodon lateral flagella

Many Molassodon genes did not have a swimming phenotype. However, many were co-expressed with lateral flagella during swimming, suggesting that the natural adaptation might encompass more than just motility, and that understanding these co-expressed genes might help us refine the natural phenotype. Two co-expressed clusters stood out: those encoding the T6SS2 Type VI secretion system (*e.g*., *tssC*) and the TonB3 active transport system (**Fig. 3A**; **Fig. S14A**).

T6SS are contact-dependent killing systems, often directed towards other bacteria. Genes encoding several T6SS2 components are differentiated in Molassodon, including structural genes, the effector RhsP, and assembly/disassembly factors (**Fig. 3F**, *top*). We reasoned that the connection between lateral flagella and antagonism might stem from the contact-dependent nature of T6SS. Motility has been shown to enhance T6SS activity in other bacteria ^26–28^, and we hypothesized that this might underlie coadaptation of these two functions in Molassodon. To provide evidence that lateral flagella-driven motility might impact the efficacy of Molassodon T6SS2, we took advantage of PVP-specific induction of *lafA* specifically in Molassodon (**Fig. 2F**) and tested T6SS2-dependent killing of prey *E. coli* under these conditions. Although killing in PVP was modest (∼0.5 log) compared to agar-based assays ^29^, deletion of *lafA* significantly increased prey survival only in Molassodon strains (**Fig. 3F**, *bottom*). While these are simplified laboratory observations, taken together with co-evolution of T6SS2 and lateral flagella and observations in other bacteria ^26–28^, these results provide experimental evidence that Molassodon T6SS2 activity could be dependent on lateral flagella in viscous conditions. This suggested to us that in a natural setting, the broader adaptation of Molassodon could be related to active antagonism.

Recent studies have also shown that T6SS-mediated killing by *Vibrio* can provide nutrients from lysed cells, and also detected a negative correlation between T6SS genes and metabolic diversity across species ^30^. We therefore investigated whether antagonism, driven by motility, could be a “hunting” strategy. Interestingly, the second co-expressed, differentiated gene cluster encodes the TonB3 nutrient uptake system (**Fig. 3G**, *top*), one of three TonB systems in *V. parahaemolyticus* (TonB1–3; *V. vulnificus* nomenclature). TonB systems transduce energy from the proton-motive force to drive active transport across the outer membrane, and recent studies have shown a functional link between T6SS, such uptake systems, and ion acquisition ^31–34^. This led us to hypothesize that coadaptation of lateral flagella, T6SS2, and TonB3 might reflect a functional linkage in the uptake of specific resources following T6SS-mediated lysis.

To test whether TonB3 could potentially facilitate nutrient acquisition from lysed bacterial prey, we grew Molassodon and Typical (n=3) wildtype and Δ*tonB3* (VP0163) strains in either rich medium (MLB) or M9 minimal salts with *E. coli* lysate as the sole nutrient source. While WT strains showed no clear growth differences under these conditions (**Fig. S15**), all Δ*tonB3* mutants reached lower densities than their respective wildtypes (**Fig. 3E**). These results suggest that TonB3 could contribute to uptake of lysate-derived nutrients in both ecospecies, although the specific substrates, whether they differ between the ecospecies, and ecological context remain to be demonstrated.

Nonetheless, together these experiments provide initial evidence for a functional link between three coadapted pathways in Molassodon—lateral flagella–driven motility, Type VI secretion, and nutrient uptake—supporting functional coherence of the genotype and providing insight for a potential ecological role.

### Convergent associations in an intermediate genotype

We noticed that some Typical *V. parahaemolyticus* strains (26% in the non-redundant dataset) also encode a *maf2* homolog (**Fig. 4A**; **Fig. S16A**; **Table S5**), encoding the flagellin glycosylation factor present in Molassodon. These “Typical *maf*+” strains were not grouped with Molassodon by GWES (**Fig. S2A**), but were detected in all geographic populations and clustered closer to Molassodon when compared using ecospecies-differentiated core genes (**Fig. S16B**). This suggested they could be an “intermediate” genotype sharing some of the same genes and variants as Molassodon.

**Figure 4.**
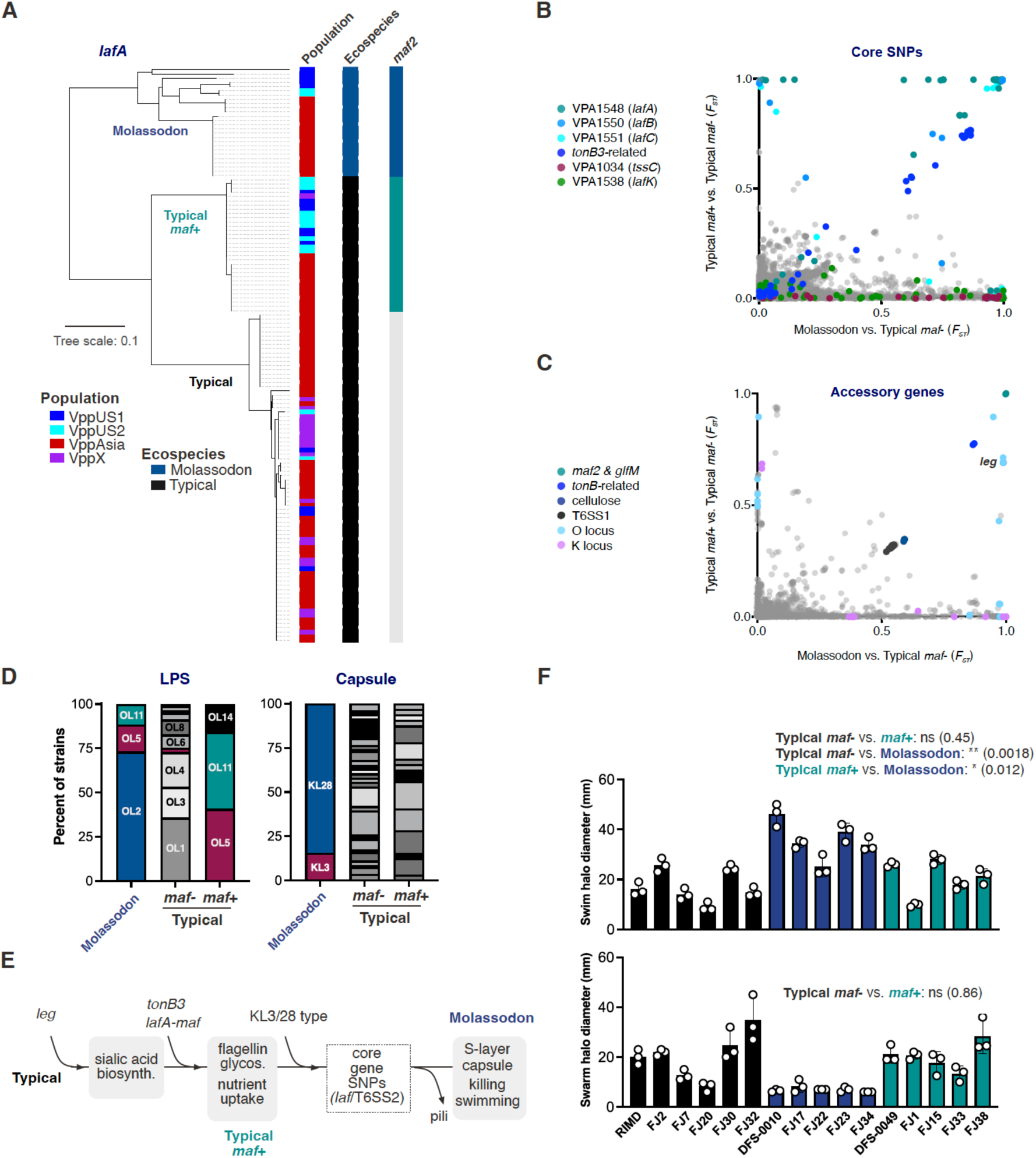
Typical *maf*+ strains recapitulate aspects of Molassodon evolution. **(A)** Neighbour-joining tree based on *lafA* for a subset of strains. Differentiated gene homologs identified by BLAST are indicated for each strain. **(B)** Core genome SNP *F_ST_* (fixation index) correlation for (Typical *maf*+) vs. (Typical *maf*-) against corresponding values for (Molassodon) vs. (Typical *maf*-). Key functions identified in this and previous ^9,10^ work are highlighted. **(C)** Correlation between *F_ST_* values for accessory genes. **(D)** Predicted serotypes for representative strains stratified into Molassodon, Typical *maf*-, and intermediate (Typical *maf*+). Key O- or K-types are indicated with colour. See also **Table S6**. **(E)** Evolutionary model for assembly of Molassodon. (**F)** Swimming and swarming of wild-type strains from different ecospecies/*maf* genotypes (Molassodon, Typical *maf*-, Typical *maf*+). ns: not significant, Student’s *t*-test.

To systematically characterize this Typical *maf*+ genotype, we performed GWAS/*F_ST_* analysis vs. Typical *maf*- strains, as well as a similar comparison with Molassodon (**Fig. S17**). Only seven core genes contained differentiating SNPs, (*e.g*., the *lafABC* genes adjacent to *maf2* and *tonB3* genes) (**Fig. S17; Table S2**). Because these loci also differ between Molassodon and Typical strains (**Fig. 1A**), we examined the correlation between *F_ST_* values for Typical *maf*+ or Molassodon strains vs. the Typical *maf*- genotype. Many SNPs in *lafABC* and *tonB* had correlated levels of differentiation for both Typical *maf*+ and Molassodon strains, although the *tonB* variants have not reached fixation in either population, since *F_ST_* values are less than 0.9 (**Fig. 4B; Table S2**). However, most SNPs were uniquely differentiated in Molassodon, such as those in other core lateral flagella (*e.g*., *lafK*) and T6SS2 (*e.g*., *tssC*) genes. As the LafK regulator has not diverged from Typical *maf*- in the intermediate strains, this suggests that distinct environmental regulation of lateral flagella may not have evolved.

Several accessory genes, such as those encoding legionaminic acid biosynthesis, also showed correlated differentiation levels between Molassodon and Typical *maf*+ strains, (**Fig. 4C**). There is a corresponding overlap in O-locus serotypes, with OL5 and OL11 found in both Molassodon and Typical *maf*+ strains (**Fig. 1E**). Both serotypes encode the *leg* genes required for Maf function. Although the third Typical *maf*+ serotype, OL14, lacks *leg* genes, all *maf*+ OL14 strains carry the KL17 capsule type, which does encode *leg* genes and is closely related to the divergent KL28 and KL3 Molassodon capsule loci. However, KL17 does not include S-layer or capsule genes related to KL28/3. Therefore, the distribution of serotypes amongst Typical *maf*+ strains further supports a functional linkage between *maf*, *lafA*, and *legI* (**Fig. 3A, E**).

We next examined the evolutionary relationship between Molassodon and the intermediate Typical *maf*+ genotype. We focused on *lafA*, which is functionally linked to *maf2* as well as being adjacent on the chromosome. Three *lafA* variants were identified in *V. parahaemolyticus*: one in Typical *maf-*strains, one restricted to Molassodon, and one in Typical *maf*+ strains (**Fig. 4A; Fig. S18A**). Moreover, we detected two *maf2* variants, with each corresponding to the Molassodon and Typical *maf*+ *lafA* alleles (**Fig. S18B**). While both *maf* versions appear to have been acquired by introgression, the Typical *maf*+ allele is closely related to a *V. antiquarius* gene and has lower diversity, consistent with more recent acquisition (**Fig. S14F, S18B**).

Divergence in *lafA-maf* between Molassodon and the “intermediate” suggested that the Typical *maf*+ genotype is not a “precursor” to Molassodon, but an independent genotype associating some of the same genes and functions. Typical *maf*+ strains also have a gene order more closely resembling that of Typical *maf*- strains, rather than Molassodon (**Fig. S3**), confirming Molassodon and the intermediate are not related clonally and are independent, convergent assemblies. However, while *maf* and *lafA* have independent origins, some shared coadapted genes appear to have a common ancestry in both populations. Phylogenetic analysis of *legI* and *tonB* revealed two distinct versions for each (**Fig. S19A, B**). Although neither showed fully fixed distribution between populations, Typical *maf*+ and Molassodon strains generally encoded the same version. Thus, while they evolved independently and have distinct lateral flagella, Typical *maf*+ strains use a part of the Molassodon gene pool, while having much less extensive coadaptation overall. For example, differentiation in *lafK* and *tssC* is clearly restricted to Molassodon (**Fig. S19C, D**).

Based on the genetic characteristics of Typical *maf*+ strains, we propose a working model for the evolution of the Molassodon genotype (**Fig. 4E**). Because legionaminic acid biosynthesis appears to be required for Maf function, acquisition of *leg* genes likely preceded *maf* acquisition. As ecological differentiation progressed, additional traits related to nutrient acquisition, predation, and regulation evolved, followed by extensive cell-surface remodeling. These changes likely reflected adaptation to altered host or environmental interactions and may have included protection from bacterial predators or phages through the S-layer, modified capsule and cellulose production, altered flagellin glycosylation, and loss of a putative pilus (VPA0713–31; **Fig. 1D**), none of which are seen in Typical *maf*+ strains.

Finally, we tested whether Typical *maf*+ strains recapitulate the Molassodon motility phenotypes. Despite sharing some differentiated loci, Typical *maf*+ strains encode distinct *maf* and *lafA* alleles and lack the more extensive functional and regulatory divergence of lateral flagella of Molassodon, such as at the regulatory gene *lafK*. Accordingly, Typical *maf*+ retained swarming ability, and did not show the enhanced swimming ability of Molassodon. However, introducing the Molassodon *maf* allele into a *maf*- *leg*+ Typical strain altered swarming (**Fig. S20**), implying that *maf* acquisition can affect motility. While Molassodon and the *maf*+ Typical strains share some differentiation, this could be an adaptation to a different niche. Nonetheless, convergent association of nutrient uptake and lateral flagella suggests that ecological differentiation in the species follows recurring axes.

## Discussion

Bacterial ecospecies are a newly recognized genotype that transcends the clonal structures most familiar to microbiologists. Here we showcase their unique features by using the Molassodon ecospecies of *Vibrio parahaemolyticus* to explore the genotype to phenotype map. We start by characterizing the Molassodon ecospecies genotype, refining our previous analysis by using a larger number of genomes, before using laboratory experiments to characterize phenotypic differences. This allows us to propose hypotheses for its ecological niche that can be tested via further laboratory and field experiments. We also identify an intermediate genotype, Typical *maf*+, which sheds light on how the genotype was assembled.

### The Molassodon genotype

Molassodon strains are in the minority in *V. parahaemolyticus*, comprising around 16.8% of environmental isolates (**Table S1**), but are found in 3 out of the 4 identifiable geographic populations and, like Typical strains contain within them nearly the entire diversity of the species in core genome SNPs and accessory genes. Molassodon and Typical strains can, however, be differentiated in three respects. Firstly, we do not find any clonal lineages that contain members of both ecospecies. Instead, Molassodon and Typical strains have coexisted on distinct clonal backgrounds for hundreds of thousands of years, such that recombination has effectively randomized the diversity within each ecospecies. Accordingly, phylogenetic methods based on undifferentiated regions of the genome produce a star-shaped tree, with many lineages of each ecospecies radiating from the same point (**Fig. 1C**), so that deeper clonal relationships are impossible to infer based on genome-wide SNP data.

Secondly, we found that gene order differentiates Molassodon and Typical strains (both *maf*- and *maf*+). The arrangement of genes on Chromosome 1 is especially distinct, and as such structural variations are more stable than shorter recombination tracts or indels, we believe this points to the age of Molassodon, with an older “clonal” frame that is still subject to rampant cross-ecospecies recombination in all regions of the genome except for differentiated genes. This also showed that differentiated regions were spread across the genome, arguing against one or two import events underlying differentiation. The position of differentiated genes away from structural variation boundaries also argues against these regions playing a role in modulating recombination frequencies between the ecospecies. Importantly, we did not observe the same distinct gene order for chromosome 1 in Typical *maf*+ strains, pointing to a newer, independent evolutionary trajectory.

Thirdly, Molassodon and Typical strains have near-fixed differences in SNPs at 48 core genes as well as in the presence or absence of over 100 accessory genes. Differences between ecospecies can accordingly be displayed as *F_ST_* plots, which are a tool widely used by animal and plant geneticists in investigating ecological differentiation. The clean, sharp peaks we obtain in our differentiation scan (**Fig. 1A**) are facilitated by our large sample of non-redundant Typical and Molassodon genomes and allow us to identify which loci are differentiated between ecospecies with high sensitivity and specificity. Indeed, the sharpness of our peaks, which often line up with gene boundaries (**Fig. S4**), compares favorably with even the most advantageous ecological genomic model systems in eukaryotes, such as river-adapted sticklebacks, where differentiation peaks typically span hundreds of kilobases.

What generates these sharp *F_ST_* peaks? The “eco” in the name ecospecies reflects the hypothesis that the peaks have been generated and maintained by natural selection and that this selection is likely to be generated by differentiation into types with distinct ecological niches. We did not find any signal of differentiation between ecospecies at most of the genome, despite large sample sizes and using fineSTRUCTURE (**Fig. S3**), a haplotype-based method that is the state of the art in fine-scale ancestry inference and that has been used for example to detect the subtle genetic differentiation between people living in different counties of England ^11,35^. This lack of differentiation in most of the genome implies that there is no meaningful barrier to recombination between ecospecies, so the differentiation needs to be maintained by some force despite constant gene flow. The only force that seems to fit the bill is natural selection. The plausibility of this explanation is further supported by preexisting annotations of the differentiated genes which are strongly skewed towards motility and cell wall biosynthesis ^10^. The fact that these variants are consistently found together on otherwise diverse genetic backgrounds represents evidence that they act together to enhance organismal fitness.

### The Molassodon phenotype

Here we present further evidence for the role of natural selection in keeping Molassodon genes together by showing that Molassodon has a coherent laboratory phenotype. This is an important intermediate goal, but our long-term purpose is to develop stronger evidence for the ecological hypothesis and to use it to zero in on laboratory conditions that are more relevant to what is happening in nature. For example, if we could identify conditions where each of the Molassodon specific variants work together to realize a fitness advantage and a second set of conditions where Typical variants work together, then we could start to be confident that our laboratory experiments were capturing the features of the natural environment that are most pertinent to ecospecies evolution. This approach can start to overcome one of the central challenges of microbiology, namely that it is difficult to assess how relevant laboratory conditions are to the real-world phenotypes we are most interested in.

To identify phenotypic differences between the ecospecies we first investigated swarming, given the large number differentiated lateral flagella genes. We found that Molassodon no longer swarmed on a surface, but all strains conserved these genes, suggesting it was not a simple loss-of-function. Instead, we found that Molassodon induces and uses its differentiated lateral flagella to swim through viscous media, where Typical strains do not. We therefore hypothesized that this swimming ability is a key feature of the Molassodon coadaptation. The complementary hypothesis is that swarming on surfaces is the key positive adaptation of Typical strains.

We took advantage of the fact that swimming is selectable in the laboratory to investigate the contribution of other Molassodon differentiated genes in addition to lateral flagella. Using Tn-seq and RNA-seq, we found that most of the genes are required and/or expressed under these conditions. Combined with mutant analysis, this revealed the first example of functional linkage among many ecospecies genes. Notably, the interaction between these genes was specific to Molassodon swimming in viscous media, suggesting we have captured some features of the natural context driving their coadaptation. While *in vitro* conditions do not necessarily recapitulate natural ecology, our data suggest that viscosity is one parameter that may differentiate the Molassodon niche, allowing us to generate hypotheses for further testing.

These experiments also give insight into the molecular basis for the relationship of Molassodon genes to the fast-swimming phenotype. Two uncharacterized genes downstream of *lafA* were required to exhibit it. Based on homology to *Shewanella* genes ^21^, we annotated them as *maf2* and *glfM*. These likely mediate sialic acid modification of the coadapted LafA flagellin. This, in turn, explained why several genes in the LPS island were also required for fast swimming: they likely encode the biosynthetic pathway for legionaminic acid, a candidate sugar substrate for Maf2/GlfM-dependent LafA modification. Consistent with this model, we observed epistasis between *lafA*, *maf2-glfM*, and *legI* during swimming. In addition to structural adaptation, differential expression of the lateral flagella is likely to play a role. In contrast, *lafA* and *legI* were not required for swimming in Typical strains. Other Molassodon genes outside the lateral flagella locus had only minor, likely indirect, effects.

The next clue that we sought to exploit from the laboratory experiments was coexpression. Molassodon lateral flagella are turned on in viscous liquids, so other differentiated Molassodon genes that are turned on simultaneously might help to explain the fitness benefit of fast swimming. The first set of genes coexpressed differentiated genes is a Type 6 secretion system, T6SS2. These systems play a role in interbacterial antagonism/competition, and we are able to show that *lafA* impacts T6SS2-mediated killing specifically in Molassodon in viscous conditions in the lab. While T6SS are often presented as competition strategies, recent genomics and experimental data has linked T6SS to nutrient acquisition ^30,31^. The differentiated *tonB3* gene is also coexpressed with lateral flagella during swimming, and we found that presence of the gene led to faster growth when the bacteria were fed on lysed *E. coli* “prey”.

These experiments provide preliminary evidence for a distinct Molassodon “hunt–kill–devour” phenotype, but could be refined in several ways. Firstly, we have tested motility, killing and nutrient uptake independently. A single experiment in which bacterial fitness is dependent on executing all three stages of the strategy could provide stronger evidence on functional linkage. Secondly, while we have shown that the T6SS2 and TonB3 are advantageous for killing and for devouring, we have not shown that the Molassodon-specific versions have a specific advantage in conditions where the strains hunt. Probing this form of allelic adaptation is likely to require a higher degree of experimental realism informed by more details about the natural Molassodon niche, especially because TonB polymorphism is not a fixed difference, suggesting that the advantage of each version is dependent on specific environmental conditions.

### Molassodon evolution

We hypothesize that Molassodon originated as a coadaptation of a small number of genes, and its function became progressively more complex and elaborate. For example, a close facsimile of the first steps in Molassodon evolution might be represented by the evolution of the *maf*+ Typical strains. These strains have evolved flagella glycosylation via coordinated changes in LafA and acquisition of the *maf* locus. These strains still swarm but are likely to have a lateral flagella-dependent motility phenotype, which we have not immediately been able to characterize. However, these convergent “intermediate” strains show similar coadaption to the *tonB* allele as in Molassodon, suggesting that motility and nutrient acquisition are intimately linked in *Vibrio* species and represent a major axis of ecological differentiation.

The *maf*+ Typical genotype is a simple coadaptation, which is likely to be easily transferrable to different genetic backgrounds. However, as the coadaptation becomes complex, selection against intermediate forms will start to restrict the mobility of particular coadaptations. There are no true intermediates between Molassodon and Typical strains, which is one reason for ecospecies designation. However, in most of the genome they are indistinguishable and for example there are Molassodon strains in three of the geographic populations, which share the allelic diversity of Typical strains from the same populations based on FineSTRUCTURE analysis (**Fig. S2B**).

Although recombination has effectively scrambled the allelic diversity of the species, deep clonal relationships can potentially be inferred through differences in gene order, some of which are not easily transferred by recombination between strains. We have found that there is a gene order configuration in Chromosome 1 that is characteristic of Molassodon strains, which is consistent with them having a deep clonal ancestor and provides further evidence that strains have not switched from Molassodon to Typical or vice versa. The gene order difference also provides further evidence that the *maf*+ Typical strains are not revertants.

### Comparison of bacterial ecospecies to other genetic structures

We now have two cleanly defined ecospecies in *H. pylori* and *V. parahaemolyticus*. The essential similarity between the two structures is shown by genome-wide *F_ST_* plots, which show regions of strong differentiation in an otherwise completely mixed genome. In both cases, ecospecies are shared between different geographic populations and there are no true intermediates. Both ecospecies are old, as indicated by genome rearrangement differences, and in *H. pylori* we have evidence that the ecospecies are at least 200,000 years old based on shared coevolution with human groups. One difference between the two ecospecies is in their evolution. In *V. parahaemolyticus* many of the differentiated Molassodon accessory genes and a few of the core genes have been imported from other *Vibrio* and related bacteria. In *H. pylori* evolution appears to have happened only by substitution and loss of key genes, likely because *H. pylori* has little access to DNA from other *Helicobacter* species within its human stomach niche.

Ecospecies are distinct from ecotypes, which have been defined theoretically as ecologically distinct, genetically cohesive units maintained by periodic selection ^36^ and operationally as phylogenetic groups with a distinct sampling distribution ^37^. Ecospecies are not identifiable as phylogenetic groups, are not periodically purged of diversity by genome-wide sweeps and are not genetically cohesive, except at ecospecies defining loci. The difference in genetic structure also influences their utility as objects of study. Because they are phylogenetically lineage with a recent origin, ecotypes are differentiated from the rest of the species throughout the genome,not just at adaptive loci, and accordingly F*_ST_* plots will be much less useful in identifying selected loci.

There are also genetic structures that share properties with ecotypes and ecospecies but do not fit into either category. For example, *Shigella* are strains of *E. coli* that are distinct enough in their phenotype that they were originally given their own species designation. An individual *Shigella* lineage can reasonably be described as an ecotype, since it is a phylogenetic group with a distinct niche. However, *Shigella* is a case of convergent evolution since similar genotypes, generated by acquisition of the same virulence plasmid, have arisen on more than one genetic background ^38^. However, recombination is substantially less effective in randomizing diversity within *E. coli* than within *V. parahaemolyticus* and accordingly, instead of being found on hundreds of distinct genetic backgrounds like Molassodon, *Shigella* occurs on only a handful. Accordingly, plots of differentiation between *Shigella* and other *E. coli* also do not produce the clean *F_ST_* plots characteristic of ecospecies (**Fig. S21**).

Ecospecies represent a progression from the early stages of adaptive differentiation that arises when strains first colonize a new niche. For example, the early-stage differentiation in a handful of loci described by Shapiro *et al.* in *V. cyclitrophicus* ^39^ resemble the early stages of differentiation such as seen in the intermediate Typical *maf*⁺ genotype that we identified and that we argue is a plausible precursor to ecospecies. Moreover, the *V. cyclitrophicus* differentiation was linked to laboratory phenotypic differences ^40^. Ecospecies will form if adaptive differentiation becomes progressively more elaborate, but with gene flow continuing to happen at a high enough rate to prohibit differentiation elsewhere in the genome. Shapiro and colleagues identified a reduction in gene flow that accompanied adaptation and emphasized its role in facilitating cohesion of the newly evolving genotypes. Thus, the species seems to be on a different trajectory, where adaptive and neutral differentiation proceeds hand in hand, leading to genome-wide differentiation that is not seen in *V. parahaemolyticus* ecospecies, despite its greater antiquity.

Ecospecies are of particular biological significance due to their high evolutionary potential, especially as precursors to speciation. Specifically, they retain the full functional flexibility of their species in most of the genome, while also developing their own recombining gene pool in ecospecies defining loci. Because they have a consistent phenotypic difference, further coadaptation will arise naturally and be shared amongst strains from the ecospecies, which is likely to slowly expand the differentiated fraction of the genome. At some point this expansion might accelerate, with selection for reduced recombination between ecospecies leading to full speciation. Ecotypes, for example, could also speciate by evolving a barrier to gene flow with other strains from their species, but this is likely to involve a genome-wide bottleneck, that will limit the immediate adaptive potential of the new species.

As objects of study, ecospecies are of particular utility because the entire complement of coadapted genes can be identified without any prior knowledge of their biological function. The approach we have followed here is thus a particularly pure example of “reverse ecology”, which first uses genomics to identify groups of strains with distinct adaptations, before attempting to infer biological and niche differences based on metadata, experiments, and/or functional annotation ^41,42^. Previous descriptive laboratory comparisons of the differentiated *V. cyclitrophicus* populations identified by Shapiro et al. ^39,40^, revealed attachment and biofilm phenotypes that are consistent with how the strains were identified (on different particles), but functional linkage to differentiated genes was not confirmed.

Here we have demonstrated how molecular genetics and global approaches such as RNA-seq and Tn-seq can be leveraged to gain insight into ecospecies function. We hypothesize that the natural phenotypic difference of Molassodon is related to hunting, killing, and devouring in viscous environments but have left field investigation to future studies. Because Molassodon and Typical strains and intermediate *maf*+ genotypes coexist (**Fig. S22**) and are detected throughout the geographical range of the species, variation in strain abundances and lateral flagella expression in different microenvironments provides a gateway for investigating the ecological niches associated with swarming and swimming behaviors, as well as their regulation in the natural environment. In the process, we can hope to understand the full ecological context of these fascinating complex adaptations.

## Materials and methods

### Genomes and sequencing data

Accessions and information for all genomes used for bioinformatics analysis are listed in **Table S1.** Complete genome sequences for some strains were generated using PacBio and Illumina sequencing at Novogene (Beijing, China) and have been deposited at NCBI under Bioproject PRJNA1235929. All other genomes are public and were recovered from NCBI. RNA-seq and Tn-seq data have also been deposited at SRA under Bioproject PRJNA1235929.

To assemble nine new genomes used for phylogenetic analysis (**Table S1**), raw reads were first quality-filtered using Filtlong (v0.2.1) ^43^, followed by genome assembly with Flye (v2.9-b1768) ^44^. Plasmid sequences were removed. A highly curated pangenome was generated for comparison of homologues by RNA-seq and Tn-seq for three Typical and three Molassodon strains: RIMD 2210633 (IPS strain, DFS-0005), SPVP110059 (DFS-0022), VP11007 (DFS-0060), VP09156 (DFS-0007), VP09285 (DFS-0008), and VP10219 (DFS-0010). A high-quality sequence for the six strains was generated using second-generation (Illumina short read) and third-generation (PacBio) sequencing at Novogene (Beijing). High quality genome assemblies were generated using Tricycler version 0.5.3 ^45^. Genomes were annotated using Prokka version 1.14.6 ^46^ and a pangenome was built using Panaroo version 1.3.3 ^47^.

### Genome-wide association studies

We conducted GWAS to identify core and accessory genes associated with Molassodon or *maf* genotypes as follows. First, we assembled a set of 1,550 public non-redundant *V. parahaemolyticus* genomes (**Table S1**). We used snp-sites ^48^ to identify single nucleotide polymorphisms (SNPs) from whole-genome alignments. Next, we calculated the SNP distance matrix and performed hierarchical clustering using the complete linkage method. A 10,000 SNP distance threshold was applied to define clusters, from which we selected the strain with the highest non-redundancy (NR) value within each cluster. To further refine the dataset, we assessed the pairwise SNP distances among the selected strains and removed those with distances below 10,000 SNPs. We then repeated this procedure using the single linkage model to extract an additional set of non-redundant strains. The final dataset comprised the union of non-redundant strains identified through both clustering approaches, ensuring a representative and diverse strain selection. We then used GWES ^10^ to define Molassodon strains (**Fig. S1**). For the *maf* GWAS based on *maf*, we defined strain “phenotype” as “*maf*+” or “*maf*-” based on detection of Molassodon *maf2* by BLAST with Molassodon *maf2*.

For the SNP-based GWAS, we employed snp-sites ^48^ to extract SNPs from whole-genome alignments, generating a variant call format (VCF) file. Association testing with either “Molassodon” or “*maf*+” was subsequently performed using pyseer ^49^, which converted inputted multiallelic SNPs into pseudo-bi-allelic categories of major allele or “other”, using default parameters. To further assess genetic differentiation between the groups, we calculated fixation index (*F_st_*) values. Weir and Cockerham’s *F_st_* ^50^ was computed using vcftools ^51^ on pseudo-haploidized bacterial SNP and accessory gene data, comparing focal strain groups (*e.g*., Molassodon) against all others. The analysis assumed clonality and negligible within-strain variation.

For the gene-based GWAS (presence/absence), we first used panaroo ^47^ to construct a pangenome and extract gene presence-absence variation from whole-genome alignments. To integrate this variation into the GWAS framework, we encoded gene presence-absence patterns as pseudo-SNP loci and subsequently employed snp-sites to generate a VCF file. The resulting VCF file was analyzed using pyseer, employing the same statistical framework as in the SNP-based GWAS. Additionally, gene-level *F_st_* values were computed to evaluate differentiation between the Molassodon and Typical (or *maf*+/*maf*- Typical) groups.

For both SNP- and gene-based GWAS, we applied appropriate quality control measures, including a minor allele frequency threshold (> 0.02), to filter out rare variants. The statistical significance of associations was evaluated using the threshold of *F_st_*≥0.5 and -log10(p) ≥10. Differentiated core genes had at least 5 SNPs with *F_st_*≥0.05 and -log10(p) ≥10, while undifferentiated genes were defined as those with zero SNPs with *F_st_*≥0.05 and -log10(p) ≥10. For the gene-based GWAS, the frequency of a gene in Molassodon strains was also required to be at least 90% or less than 10%.

### Identification of orthologs in *V. parahaemolyticus* and other species in KEGG

The sequences from Molassodon strain UCM-V493 (vph) or Typical *maf*+ strain VP08404 were used for blastn at KEGG and the workflow mafft_default-none-none-fasttree_default as described. Trees were visualized with iTOL ^52^. KEGG genomes are listed in **Table S1**. Introgression events were defined as those where the Molassodon allele branched away from Typical *V. parahaemolyticus* strains based on analysis of KEGG homologues.

### Phylogenetic analysis

To generate phylogenetic trees in Fig. 1C, a set of publicly available genomes plus nine newly-sequenced genomes (see above) were used, selected to include strains available in our lab for experiments as well as representatives from all groups of interest (*maf* genotype, ecospecies, population) (**Table S1**). To generate the *lafA* (VPA1548 in RIMD2210633) tree, the nucleotide sequence from UCM-V493 (VPUCM_21376) was aligned against the 139 strains using BLASTN. This revealed over 82% coverage of the VPA1548 (*lafA*) region in the RIMD 2210633 reference genome, which was subsequently used for phylogenetic reconstruction. In total, 124 SNPs were identified in this region and were used to construct a Neighbor-Joining (NJ) tree using TreeBest. To generate trees based on differentiated/undifferentiated genes, we extracted all SNPs within these regions and constructed an Approximate Maximum Likelihood Tree phylogenetic tree, again using FastTree ^53^. Trees were visualized using iTOL ^52^. To generate the phylogenetic tree in **Fig. S1A**, each genome was aligned to the RIMD2210633 reference using NUCmer with the --mum option. Alignments were filtered using delta-filter -1 to retain one-to-one best matches, followed by additional stringent filtering. Aligned regions were extracted in reference coordinates, non-ATCG characters were converted to gaps, and low-coverage genomes were excluded prior to SNP extraction. SNPs were subsequently identified from the filtered alignments, and repeat-region SNPs were removed before downstream analyses and phylogenetic reconstruction.

### Analysis core genome *F_st_* in populations, deep clones, and *E. coli*/*Shigella*

To identify clonal lineages among the 1,550 non-redundant *V. parahaemolyticus* strains, we first calculated a pairwise SNP distance matrix and conducted hierarchical clustering using the single linkage method. Clonal groups were defined by cutting the resulting dendrogram at SNP distance thresholds of 45 kb and 50 kb. Lower thresholds identify smaller groups, while larger cutoffs are likely to cluster unrelated strains, based on the median SNP distance of 54,984 within the VppAsia population. The largest putative clonal groups within the VppAsia population obtained under each threshold consisted of 18 (45 kb) and 21 (50 kb) strains, respectively, and the pairwise distribution for each group was clearly distinct from that observed for unrelated strains (**Fig. S5A)**.

To control for sample size effects, we randomly selected 18 and 21 strains from the VppAsia Molassodon lineage to match the clonal groups. For each case, a background population of 950 non-Molassodon VppAsia strains was used to minimize confounding due to broader population structure.

We conducted both genome-wide association studies (GWAS) and *F_ST_* analyses on the following four pairwise comparisons:

1. 18-strain clonal group (45 kb threshold) vs. 950 non-Molassodon VppAsia strains
2. 21-strain clonal group (50 kb threshold) vs. 950 non-Molassodon VppAsia strains
3. 18 randomly-selected VppAsia Molassodon strains vs. 950 non-Molassodon VppAsia strains
4. 21 randomly-selected VppAsia Molassodon strains vs. 950 non-Molassodon VppAsia strains

Each pair was treated as a binary phenotype in GWAS and as two distinct populations for *F_st_* estimation.

To assess broader patterns of population differentiation and to provide a comparative framework for the Molassodon-based analysis, we constructed two additional datasets for population-level *F_st_* estimation:

1. VppAsia (n = 1,209) vs. VppX (n = 176)
2. VppAsia (n = 1,209) vs. VppUS (n = 165; combining both VppUS1 and VppUS2).

To compare core genome differentiation between *E. coli* and *Shigella* isolates, a previously published non-redundant dataset was used (n=500) ^54^.

In each case, *F_st_* was calculated between the two groups to quantify genetic differentiation across population lineages.

### Recombination-Based Age Estimation

To estimate the ages of the deep clonal groups and the Molassodon lineage, we used a direct recombination-clock approach based on pairwise core-genome SNP distances. This approach follows the recombination-based divergence framework proposed by Cui et al. ^9^ but estimates divergence times directly from pairwise SNP distances without fitting an *N_e_r* model.

Assuming that sequence divergence between two strains accumulates primarily through homologous recombination, the probability that a genomic site remains unrecombined since the two strains shared a common ancestor at time *T* is:

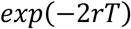

Thus, the expected divergence between two strains can be expressed as:

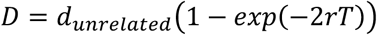

where *D* is the observed core-genome SNP distance between two strains, *d_unrelated_* is the median SNP distance between unrelated strains, and *r* is the recombination rate per site per year. Rearranging this equation gives:

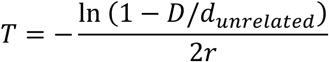

To restrict the analysis to a single population, we used VppAsia strains to define the background divergence level. The median pairwise distance among unrelated VppAsia strains was *d_unrelated_* = 54,875 SNPs. We adopted a recombination rate of *r* = 1.7 × 10^12^ per site per year, as previously estimated for *V. parahaemolyticus* populations.

For each group, pairwise divergence times were calculated for all strain pairs using the equation above. We then calculated (i) the mean pairwise SNP distance (*D_mean_*), (ii) the mean unrecombined genome fraction

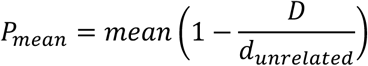

and (iii) the mean pairwise age

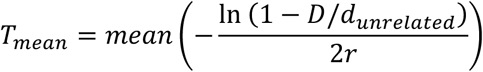

These quantities provide intuitive summaries of the typical genetic divergence, the fraction of the genome remaining unrecombined, and the corresponding divergence time within each group. The analysis was applied to an 18-strain deep clone defined using a 45 kb SNP-distance threshold, a 21-strain deep clone defined using a 50 kb SNP-distance threshold, and the 220-strain Molassodon lineage within the VppAsia population.

### Additional bioinformatic analysis

*In silico* serotyping was performed using the *V. parahaemolyticus* Kaptive database ^12,13^. Synteny analysis at single loci (*e.g*., the *slr4* region) was performed with webflags (www.webflags.se) ^55^. To perform core genome synteny block analysis, SibeliaZ ^56^ was used for multiple sequence alignment and to build local collinear blocks, and then maf2synteny ^57^ was used to merge small local collinear blocks into larger synteny blocks. Structures of Slr4 homologues and protein complexes (Maf2 + GlfM) were predicted with AlphaFold2 or Alphafold multimer at Colabfold ^22,23^. Structures were visualized with Pymol version 3.0.2. RNA-seq coverage was displayed with Integrated Genome Browser ^58^. The non-redundant *V. parahaemolyticus* dataset (n=1550) was analyzed using FineSTRUCTURE ^11^ as described previously ^9,10^.

### Bacterial lab strains and culture conditions

Bacterial strains are listed in (**Table S7**). Most *V. parahaemolyticus* wild-type lab strains were obtained from the Shenzhen Center of Disease Control and Prevention, except for RIMD 2210633 reference isolates, which were kind gifts from Yiquan Zhang (Affiliated Hospital 3 of Nantong University) and Qiyao Wang, East China University of Science and Technology. A set of sympatric natural strains was isolated from seawater from the coast of Fujian Province, China (平潭, Pingtan and 泉州, Quanzhou) using CHROMagar™Vibrio plates according to the manufacturer’s recommendations. *V. parahaemolyticus* strains were routinely grown either on Lysogeny broth (LB: 1% (w/v) tryptone, 0.5% yeast extract, 1% NaCl, Sangon) plates or MLB (LB with 3% NaCl final concentration) plates at 37°C/30°C, respectively, with 1.5% agar or in broth culture in LB/MLB with shaking at 200 rpm at 37°C/30°C in an aerobic atmosphere. Where appropriate, media was supplemented with marker-selective antibiotics (5/25 μg/ml chloramphenicol (Cm) for *V. parahaemolyticus*/*E. coli*, 50 μg/ml kanamycin (Kan)). *E. coli* strains (**Table S7**) were grown aerobically at 37°C in LB broth or on LB agar supplemented with the appropriate antibiotics for marker selection as listed above.

### Motility assays

Strains were grown in MLB until log phase, and then 0.05 OD_600_ was inoculated into swim or swarm plates. Swim plates were made from 0.25-0.325% agar as described previously ^17^. For swarming, we used either previously described BHI agar (1.5%) ^17^, or for Tn-seq, LB agar with 2% NaCl. Plates were incubated at 30°C for approximately 16 hours until halos were observed. For diameter measurements, if the colony was not round, which means the longest diameter is more than 1 mm longer than the shortest diameter, we took the average between the longest and the shortest diameters.

### General recombinant DNA techniques

All plasmids generated and/or used in this study are listed in **Table S14**. Oligonucleotide primers (Sangon) are listed in **Table S15**. DNA constructs and mutations were confirmed by Sanger sequencing (Sangon). Restriction enzymes, *Taq* polymerase for validation PCR, and T4 DNA ligase were purchased from Tiangen, ThermoFisher Scientific, or New England Biologs. For cloning purposes, Phanta high-fidelity DNA polymerase was used (Vazyme). Some plasmids were generated with the Sangon Seamless cloning kit.

### *V. parahaemolyticus* deletion mutant construction

*V. parahaemolyticus* deletion mutant strains were generated via recombination with suicide plasmids generated based on pDM4 ^59^, as described previously ^17,60^. As an example, the deletion of *lafK* in RIMD 2210633 is described. Oligonucleotides and plasmids used for other mutants are listed in **Tables S14** & **S15**. Approximately 1kb upstream/downstream of the region to be deleted was amplified with primers DFO-0020/0021 and DFO-0022/0023 from WT genomic DNA (DFS-0191), respectively. This introduced a short (approximately 25 bp) region of homology between the two 1kb fragments for overlap PCR, as well as *Xba*I and *Xho*I restriction sites for cloning into pDM4. The upstream and downstream fragments were mixed in an equimolar ratio and subjected to overlap PCR using DFO-0020/0023, and pDM4 was amplified with DFO-0018/0019 and digested with *Dpn*I to remove template DNA. The overlap PCR product and pDM4 were digested with *Xba*I/*Xho*I and then ligated overnight with T4 DNA ligase.

In some cases, the Sangon seamless cloning kit was used. Here, PCR primers for upstream and downstream homology regions also introduced regions overlapping pDM4 (amplified with DFO-0412/0413). For example, to generate a *lafK* deletion construct for Molassodon strains, fragments were amplified with DFO-0410/0281 and DFO-0282/0411. Fragments were mixed in approximately equimolar ratios and fused after addition of seamless cloning mix for 30 min at 50°C.

Ligations/seamless cloning reactions were transformed into DH5α λ*pir* and Cm^R^ colonies were verified by colony PCR with DFO-0044/0045. Verified plasmids (*e.g*., pSSvSH3 for *lafK*) were transformed into *E. coli* SM10 λ*pir* or *E. coli* MFD λ*pir,* followed by conjugation into the *appropriate V. parahaemolyticus* recipient strain. Transconjugants with single crossovers were first selected on MMM (minimal marine medium) plates with 10 µg/ml Cm. Single colonies were restreaked to purity and then transferred to MMM plates containing 15% sucrose to counter-select for double-crossover strains. Deletion clones were verified by colony PCR with DFO-0024/0023 (for Δ*lafK*) and checked for Cm sensitivity to ensure the second recombination had occurred.

### Expression of maf2-glfM in Typical V. parahaemolyticus

To overexpress Molassodon *maf2-glfM* in Typical strains, the two genes with their native RBS were amplified with short, overlapping sequences to the pMMB207 plasmid using DFO-0942/0943 from DFS-0010 gDNA. The pMMB207 plasmid backbone carrying the Ptac promoter was amplified by PCR with DFO-0437/0438. The two l fragments were then assembled into a plasmid by ligation-independent cloning (CPEC, circular polymerase extension cloning) ^61^ and transformed into DH5α λ*pir*. The resulting plasmid, pWHQ222, was verified by colony PCR with DFO-0306/0187 and sequencing with DFO-0168, and then conjugated from *E. coli* MFD λ*pir* into the appropriate recipient *V. parahaemolyticus* strain.

### sfGFP reporter construction

To generate a transcriptional reporter for the Molassodon *lafA* (DFS0010_04645) promoter, approximately 120 bp upstream of the *lafA* start codon was amplified with DFO-0300/0302 from DFS-0007 gDNA to add an *Xba*I site at the 5’ end and a region overlapping the strong non-native ribosome binding site (RBS) and GFP from pXG10-SF ^62^ at the 3’ end. The superfolder GFP (sfGFP) gene was amplified with the RBS from pXG10-SF with DFO-0180/0139, which introduced a *Pst*I site at the 3’ end of sfGFP. The two fragments were fused by overlap-extension PCR with DFO-0300/0139. The conjugatable replicating plasmid pMMB207 ^63^ was amplified by inverse PCR with DFO-0279/0185 to add *Xba*I and *Pst*I sites and digested with *Dpn*I to remove template DNA. The vector and overlap PCR fragments were then digested with *Xba*I and *Pst*I, ligated, and transformed into DH5α λ*pir*. The resulting plasmid, pSSvSH47, was verified by colony PCR with DFO-0306/0187 and sequencing with DFO-0168, and then conjugated from SM10 λ*pir* into the appropriate recipient *V. parahaemolyticus* strain.

To generate the corresponding reporter for RIMD2210633 P*lafA*, CPEC was employed to replace the Molassodon *lafA* promoter in pSSvSH47. Approximately 170 bp upstream of the *lafA* start codon with short, overlapping sequences was amplified with DFO-0384/0443 from RIMD2210633 gDNA and a GFP reporter skeleton was amplified with DFO-0442/0385 from pSSvSH47. The two linear DNA fragments were then assembled into plasmids using a Ready-to-Use Seamless Cloning Kit (Sangon) and transformed into DH5α λ*pir*. The resulting plasmid, pWHQ201, was verified by colony PCR with DFO-0306/0187 and sequencing with DFO-0168, and then conjugated from SM10 λ*pir* into the appropriate recipient *V. parahaemolyticus* strain.

### P*lafA* reporter assays

To measure the P*lafA* reporters, strains were grown in MLB with or without 3% (w/v) PVP (polyvinylpyrrolidone K90, Xushuo Biotechnology Co., Ltd., Shanghai, China) at 30°C, and then optical density (OD_600_) and GFP (excitation 485 nm/emission 528 nm), were measured every 15 minutes.

### RNA isolation and cDNA library preparation for RNA-seq

Bacterial cells from cultures or motility plates were mixed with 0.2 volumes stop mix (5% phenol, 95% ethanol) and immediately frozen at -80°C. RNA was extracted from thawed cell pellets for total RNA using hot phenol, respectively, as described previously ^64^. Total RNA was digested with DNase I, depleted of rRNA, and converted to cDNA libraries at Novogene, Inc (Beijing, China).

### Processing of RNA-seq data and RNA-seq expression analysis

Fastp version 0.23.4 was used for quality control of raw reads with default parameters ^65^. Reads were then mapped to the corresponding reference genome using Segemehl version 0.3.4 with default parameters ^66^. FeatureCounts version 2.0.3 was used for gene expression quantification with parameters “-B -C -f - p -s 1 -O --fraction --countReadPairs --ignoreDup –primary” ^67^. Differential expression was performed using R package DESeq2 version 1.42.1 using the output from featureCounts without normalization ^68^. Gene set enrichment analysis was performed using R package clusterProfiler version 4.10.1 on differentially expressed genes passing a filter of adjusted P-value < 0.05 ^69^. Gene set classifications include GO terms ^70^, KEGG pathways, KEGG KO ^71^, COG categories ^72^, and Molassodon genes based on GWAS. Other general sequencing file and alignment file manipulations (*e.g*., read reverse complementation, alignment statistics, genome coverage file generations, etc.) and calculations (e.g. TPM normalization) were performed using Samtools version 1.19 ^73^ and Julia. Plots were generated using Julia package Makie.jl ^74^.

### Tn-seq data generation

To generate a high-density Tn pool, we used a Mariner transposon delivered by conjugation on a suicide (R6K origin) plasmid. We modified the plasmid pCAT-Mariner^75^ to be compatible with current Illumina platforms as follows. Specifically, Illumina sequencing components were added to both sides of the Mariner transposon repeat region through PCR using primers DFO-690/672, DFO-690/0673, DFO-0689/0684, and DFO-689/683 from pCAT-Mariner. The four resulting linear DNA fragments were then assembled into plasmids CPEC and transformed into DH5α λ*pir*. The resulting plasmid, pWHQ011, was verified by sequencing with primers DFO-672 and DFO-673 for each side of the repeat region.

Following conjugation with *E. coli* MFD λ*pir*, putative Tn mutants were selected on LB with Cm at 10 µg/ml. Approximately ∼500,000/∼800,000 colonies (Typical/RIMD 2210633/DFS-0191 and Molassodon/DFS-0010) were recovered and immediately frozen at -80°C with 25% glycerol. After sequencing (see next section) we detected 168,777 for the RIMD 2210633 pool and 331,277 unique insertions for the DFS-0010 pool.

Motility selection was applied as follows. For swarming (RIMD 2210633 only) cells were recovered from the -80°C freezer by shaking in MLB Broth with Cm at 10 µg/ml for 3 hours at 30°C. Cells were then resuspended in MLB and approximately 10^7^ CFU in 10 µl was spotted on 2% MLB plates, which we found generated robust swarming, 10 times. After approximately 16 hours of incubation at 30°C, we harvested the outermost edge of swarm halos (Swarm-out), as well as the inside (Swarm-in) as a reference. For swimming (both Typical and Molassodon), cells were recovered from the -80°C freezer by shaking in MLB Broth with Cm at 10 µg/ml for 3 hours at 30°C. Next, approximately 10^8^ CFU in 100 µl was inoculated into swim agar ^17^ in a large beaker to ensure enough material could be recovered for library preparation. Assays were incubated at room temperature for 16 hours. As a reference, we inoculated the same swim agar with the mutant pool and shook the tube at room temperature during the same time period to remove the selection for motility, maintain the same nutrient conditions, and obtain enough material for sequencing. To isolate mutants from the swimming agar, the swimming agar containing mutants was collected and centrifuged at 3500 rpm for 10 minutes to precipitate the bacterial cells. Subsequently, the bacteria were repeatedly washed with an equal volume of PBS until the agar was completely removed from the precipitate.

Tn-seq libraries were generated from input (original mutant pools, no selection), reference, and motility libraries, essentially as described previously ^76^, as follows. Genomic DNA was extracted using the FastPure Blood/Cell/Tissue/Bacteria DNA Isolation Mini Kit (Vazyme). Subsequently, 5 µg of DNA was digested with *Mme*I (New England Biolabs), and transposon-sized fragments of approximately 1500 bp were extracted via gel extraction (Gel DNA Recovery Kit, Zymo) after separation on a 2% agarose gel. Annealed sequencing adapters were ligated overnight at room temperature. The ligated samples were then used as templates for final library PCR enrichment, which was performed for 22 cycles using an adapter-specific primer (DFO-0762) and a transposon-specific primer containing different Illumina indices (DFO-0936 - DFO-0941). The resulting 171 bp products were purified by gel extraction and sequenced on the Illumina NovaSeq 6000 system.

### Tn-seq data analysis

Raw data were first subjected to quality control using fastP (v0.23.4) and processed with cutadapt (v4.8) to remove sequencing adaptors and transposon sequences. The remaining 14 - 16 bp transposon-junction sequences were mapped to RIMD 2210633 or DFS-0010 chromosomes using Bowtie (v1.3.1) ^77^. Subsequently, the read counts of each insertion site were tallied using the TraDIS toolkit ^78^. To compare the changes in the relative abundance of insertions in each gene across different samples, the total read counts of each sample were normalized to the lowest count value in each group. Following normalization, the log - ratio of output read counts to reference read counts was calculated for each gene.

### Transmission electron microscopy

For visualization of lateral flagella, the appropriate strains were grown in MLB liquid culture or on MLB plates. For liquid cultures, 100 µl of overnight culture was harvested, gently washed twice with 1x PBS, and resuspended in 1x PBS to a final OD600 of approximately 0.1. For samples from agar plates, we resuspended part of the colony in 1 ml of 1x PBS. A 300 mesh copper grid (EMCN) was prepared by glow discharge (PELCO easiGlow 91000), and 10 µl of washed cells was added. After 1 min, liquid was removed gently with a filter paper and the grid was washed with 10 µl 1x PBS. The grid was then negative-stained with uranyl acetate for 20 seconds. The uranyl acetate was removed with a filter paper. The grid was quickly washed once with sdH_2_O and dried under a heat lamp.

### Type VI killing assay

T6SS2 killing activity was measured against *E. coli* BW25113 (DFS-2143) carrying a Kan^R^ plasmid (pWHQRed2) to allow plating for colony-forming units (CFUs). To generate the plasmid, approximately 450 bp upstream of the *gyrB* start codon with short, overlapping sequences was amplified with DFO-1078/1072 from RIMD 2210633 gDNA, the RFP gene was amplified with its RBS from pRP0122-Pbe-AmCyan_Bc3-5_turboRFP with DFO-1073/1087 and a plasmid backbone was amplified with DFO-1088/1081 from pET28a. The three linear DNA fragments were then assembled into plasmids using a Ready-to-Use Seamless Cloning Kit (Sangon) and transformed into DH5α λ*pir*. The resulting plasmid, pWHQRed2, was verified by colony PCR with DFO-1078/0187 and sequencing with DFO-0187, and then electroporated into the *E. coli* BW25113 strain. Predator strains included Typical and Molassodon wild-type, Δ*hcp2*, and Δ*lafA* strains (**Table S7**). To initiate the killing assay, overnight cultures of prey and predator strains were washed with PBS one time and resuspended by pipetting. Predator and prey suspensions were diluted to a final OD_600_ of 0.5 with M9 broth (Sangon) with 2% (w/v) NaCl (final concentration) and 5% PVP. Predator and prey strains were mixed at a ratio of 10:1 and incubated at 30°C. A control with *E. coli* only was included. After 3.5 hours, serial dilutions were spotted onto antibiotic-containing LB agar plates to enumerate prey colony forming units (CFU/ml). Data were normalized using the CFU of the corresponding single *E. coli* strain group.

### Growth assays

First, a whole cell lysate of prey *E. coli* BW25113 was generated as follows. Approximately 60 OD_600_ of cells were harvested from LB broth cultures grown at 30°C until log phase and washed three times with PBS and lysed by sonication. The sonicated lysate was filtered through a 0.22 μm filter (Sangon). *V. parahaemolyticus* strains were grown in pre-culture in 2 ml MLB at 30°C until approximately log phase, washed three times with 1× PBS, and inoculated into test media at a final OD_600_ of 0.5. Each strain was grown in triplicate in 200 µl of MLB or M9 (salts only + NaCl to 2%; no carbon source) with ⅔ volume *E. coli* lysate with shaking at 30°C. Results shown are representative of two independent experiments.

### Total protein analysis

Protein expression in *V. parahaemolyticus* was performed by SDS-PAGE of whole-cell lysates. Strains were grown in MLB at 30°C until log phase, and then cells were harvested by centrifugation at 13,000 rpm. Cell pellets were resuspended in 100 μl of 1× protein loading buffer (62.5 mM Tris-HCl, pH 6.8, 100 mM dithiothreitol, 10% (v/v) glycerol, 2% (w/v) SDS, 0.01% (w/v) bromophenol blue) and boiled for 8 min. Next, 0.1 OD_600_ of cells were loaded per lane on 12% SDS-polyacrylamide gels. After electrophoresis, protein bands were visualized by staining with Coomassie Blue (Sangon).

### Statistical analysis

Statistical tests for experimental data were performed using Graphpad prism.

## Acknowledgements

We thank Qiyao Wang, Yiquan Zhang, Kim Orth, and Melanie Blokesch for providing strains and advice, and Elise Tourrette for assistance with GWAS.

## Author contributions

Performed experiments: SLS, ZJ, HW, ZX. Bioinformatic analysis: SLS, ZJ, HW, HF, WX, CY. Designed research: SLS, DF. Materials and strains: YC. Analysis and interpretation: SLS, DF. Manuscript writing: SLS, DF.

## Data and materials availability

Newly-sequenced genomes, RNA-seq and Tn-seq data have been deposited at SRA under Bioproject PRJNA1235929.

