## Supplementary Material for "Functional coherence of bacterial ecospecies"

This document contains:
**Supplementary Figures S1-S22**
**Supplementary References**

### Supplementary Figures

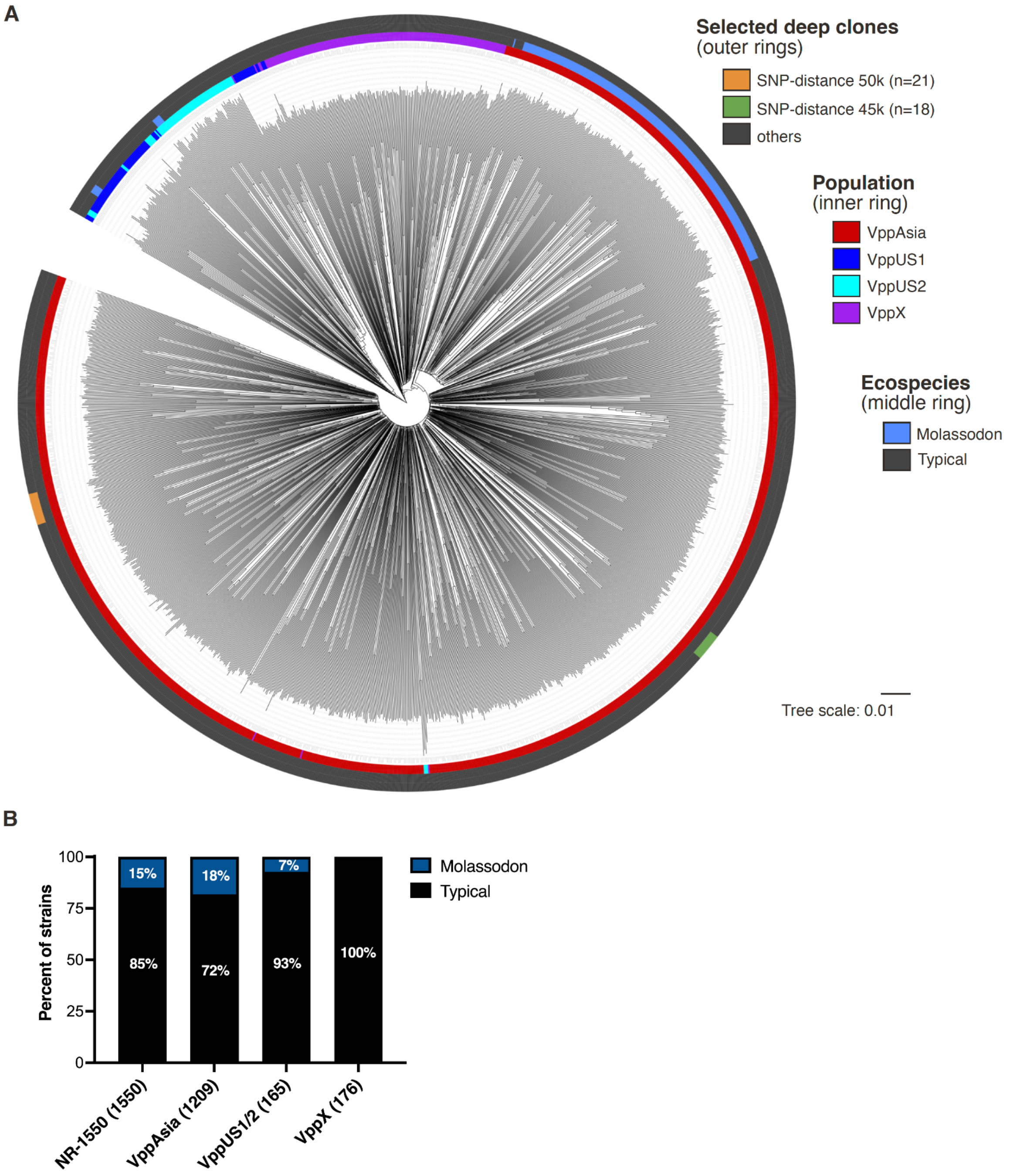


**Supplementary Figure S1. Non-redundant dataset overview. (A)** Neighbour-joining tree of non-redundant dataset inferred from a filtered SNP-only alignment. Geographic population, ecospecies, and selected clones indicated. Branch lengths represent substitutions per retained SNP (see **Methods** for details). **(B)** Proportion of Molassodon and Typical strains in the non-redundant (NR) dataset and populations. VppUS: VppUS1 + VppUS2.

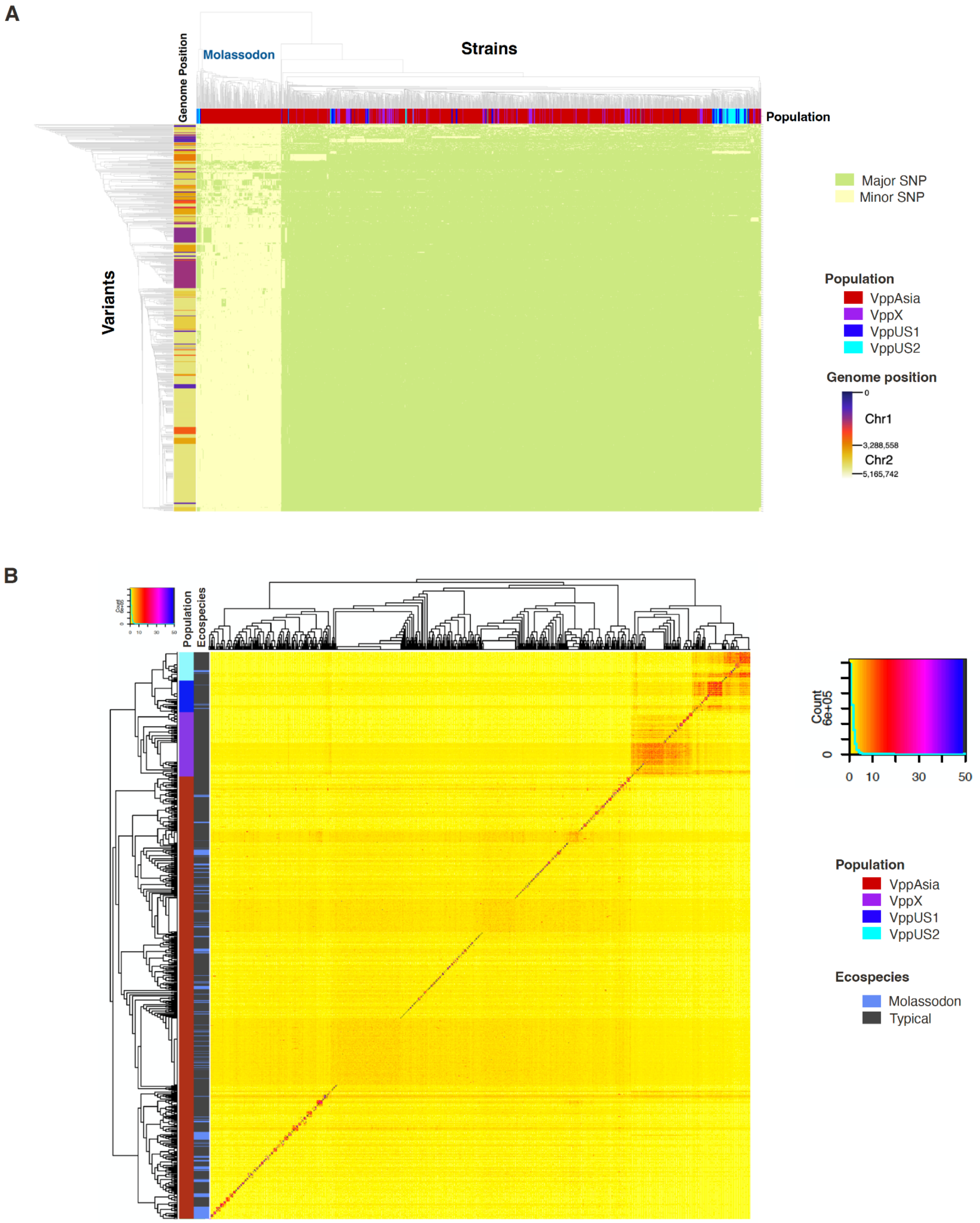


**Supplementary Figure S2. Population structure analysis of the non-redundant *V. parahaemolyticus* dataset. (A)** Genome-wide epistasis scan of a non-redundant V. parahaemolyticus dataset (n=1550). **(B)** FineSTRUCTURE analysis of the *V. parahaemolyticus* non-redundant dataset.

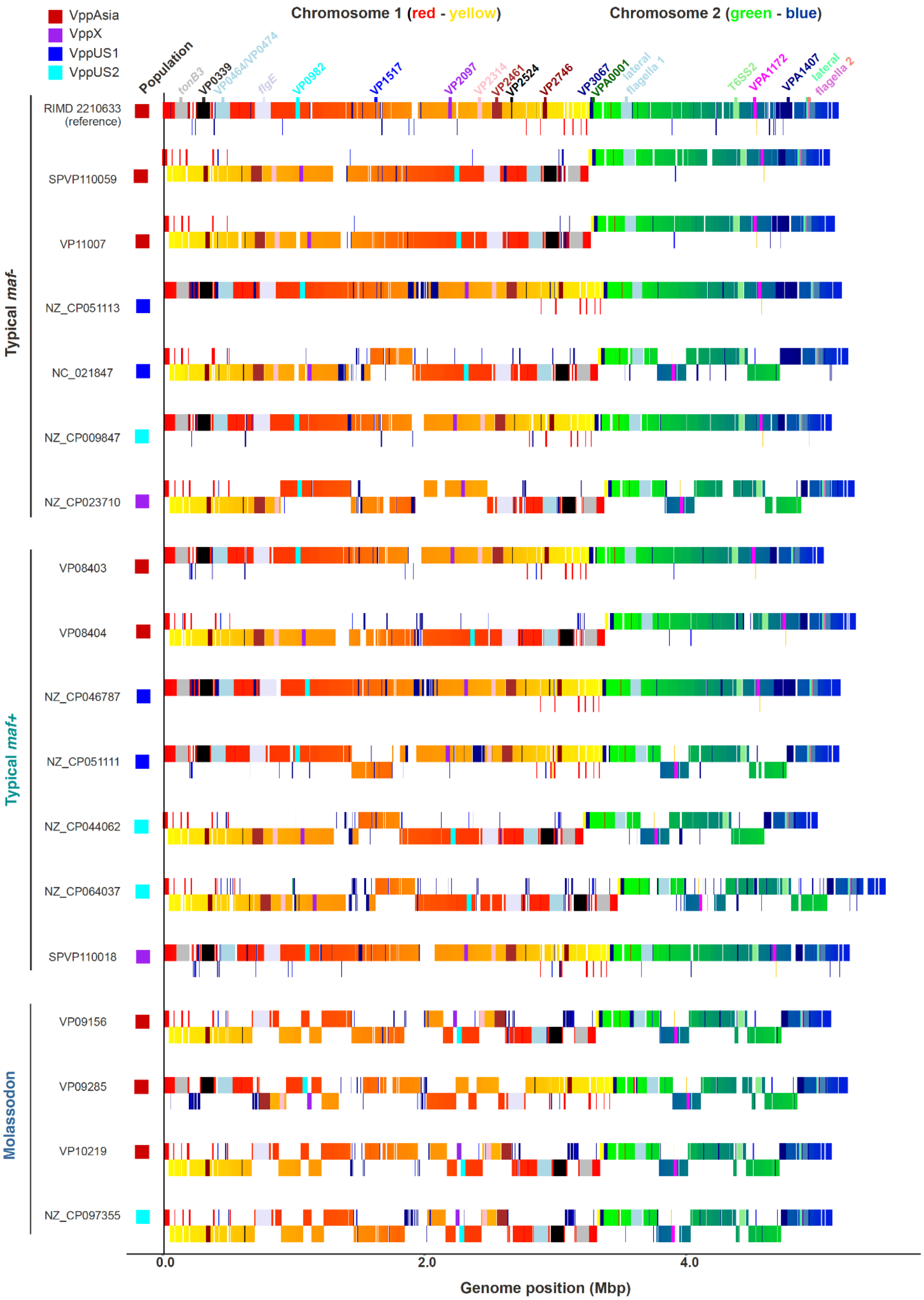


**Supplementary Figure S3. Synteny block analysis.** Order of synteny blocks on Chromosomes 1 and 2 for Typical *maf-*/*maf+* and Molassodon strains. Blocks are coloured according to the order for the RIMD 2210633 (Typical) reference strain. Selected available complete genomes were included in the analysis.

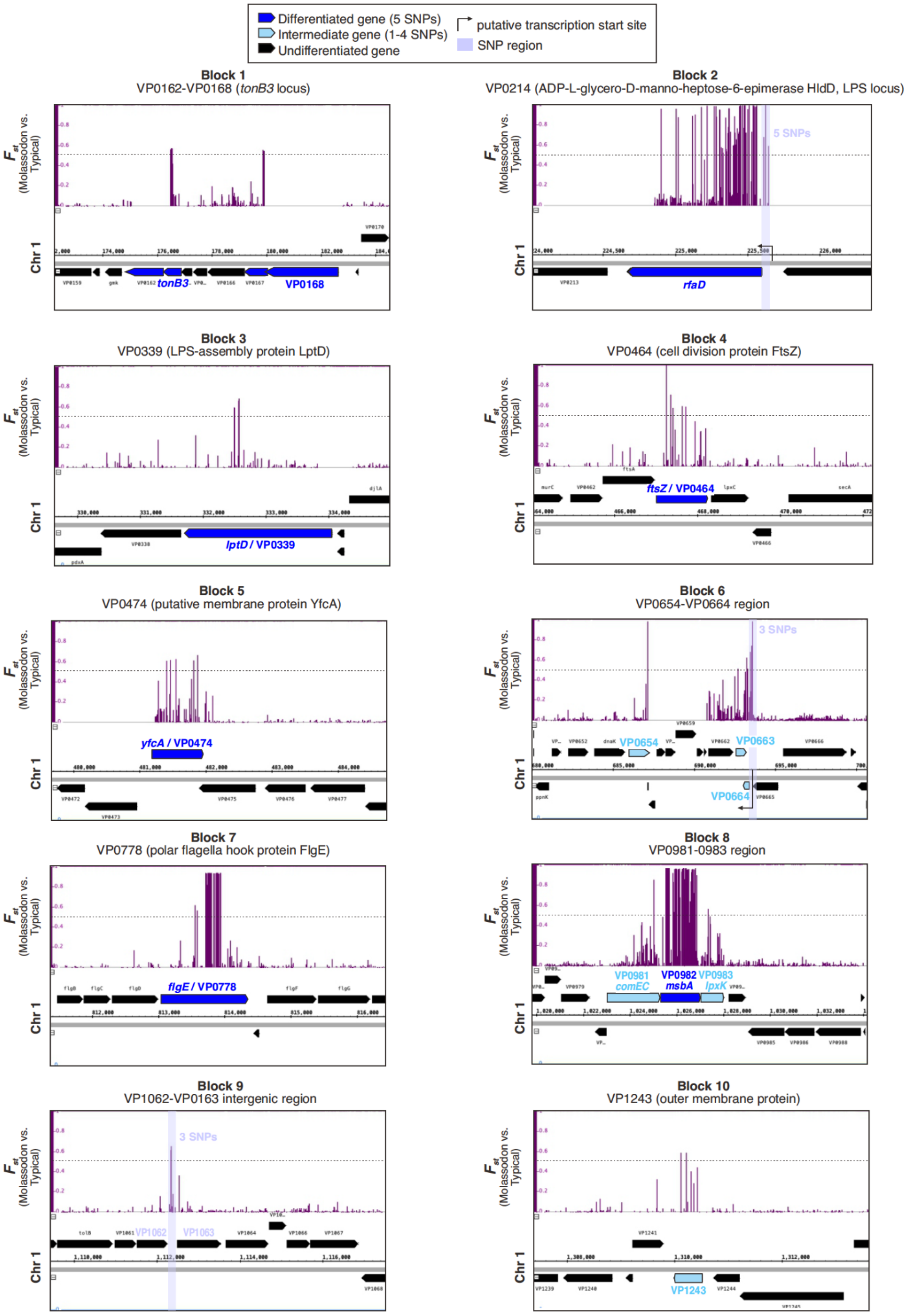


**Supplementary Figure S4. Zoom-in on $F_{ST}$ values for reference genome blocks with at least one differentiated SNP between Molassodon vs. Typical.** See also **Table S4.**

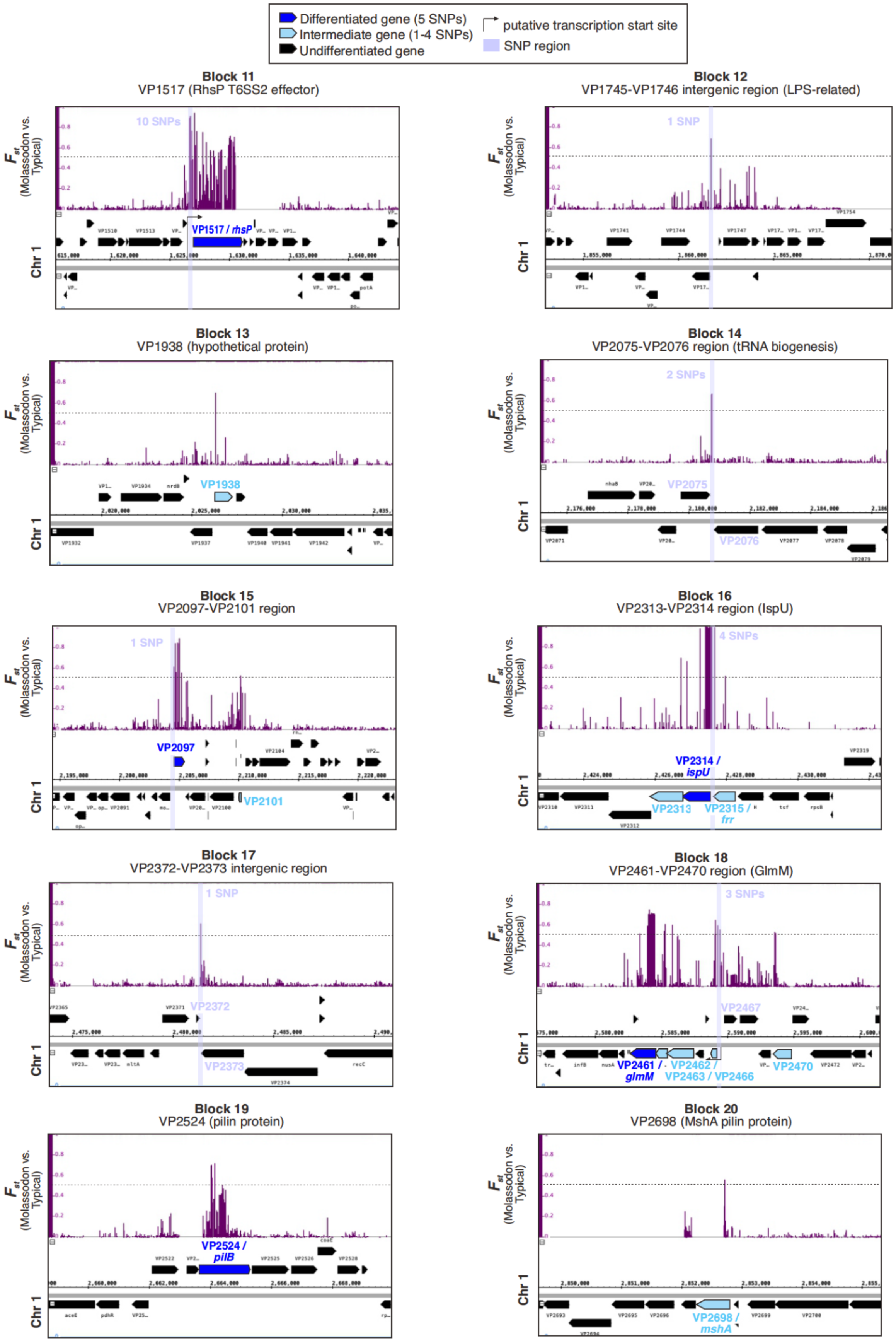


**Supplementary Figure S4 (continued). Zoom-in on $F_{ST}$ values for reference genome blocks with at least one differentiated SNP between Molassodon vs. Typical.** See also **Table S4.**

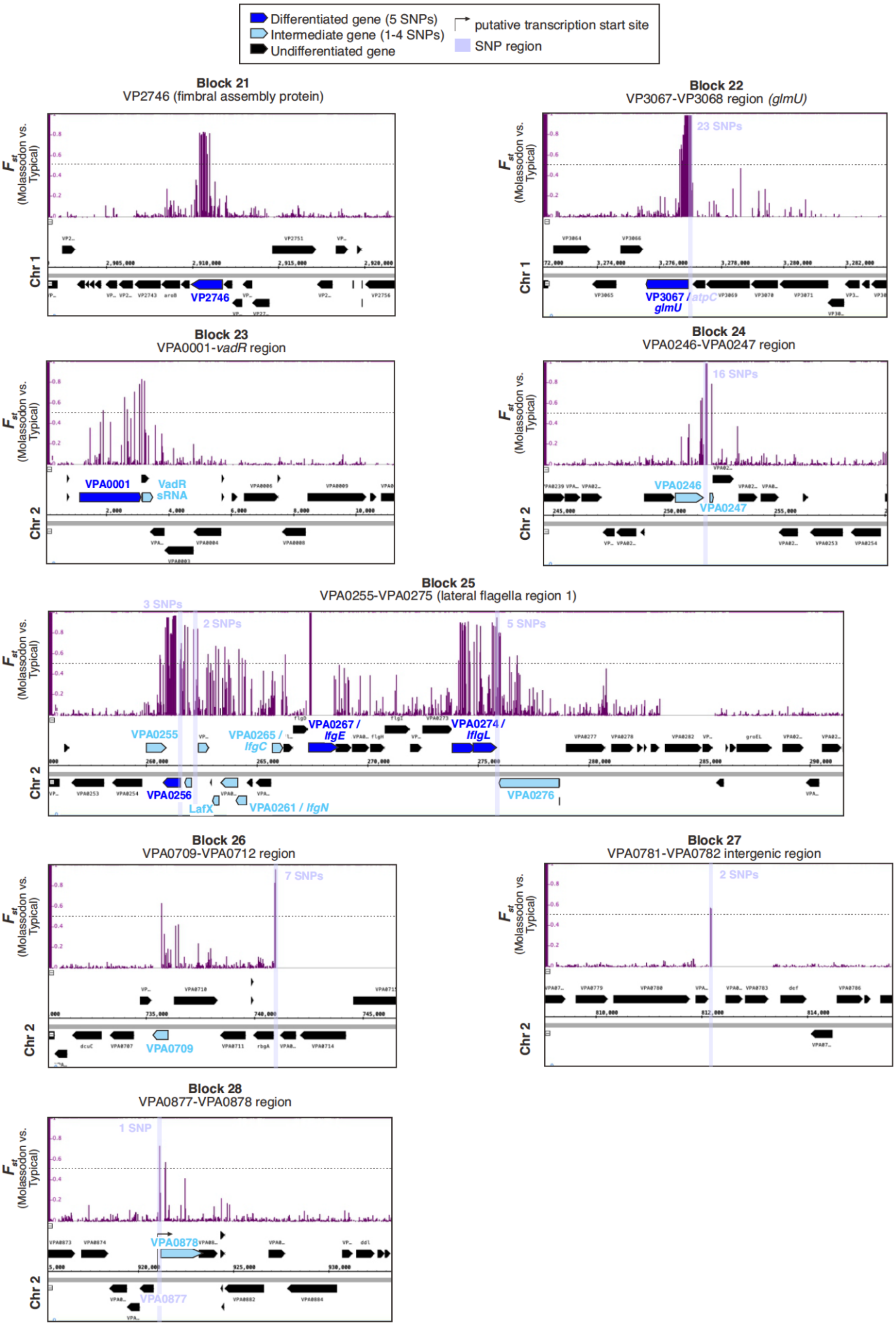


**Supplementary Figure S4. Zoom-in on *F*$_{ST}$ values for reference genome blocks with at least one differentiated SNP between Molassodon vs. Typical.** See also **Table S4.**

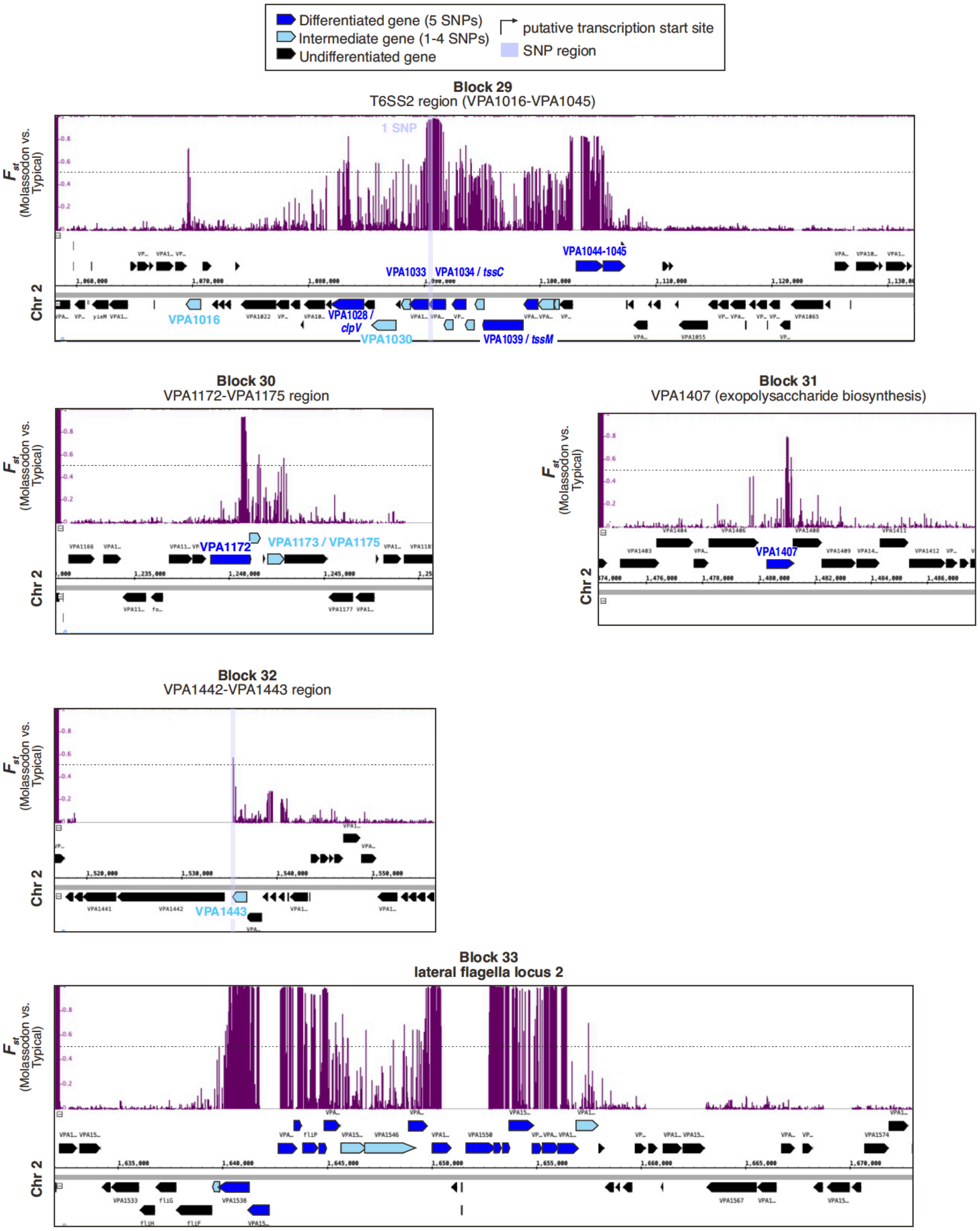


**Supplementary Figure S4. Zoom-in on $F_{ST}$ values for reference genome blocks with at least one differentiated SNP between Molassodon vs. Typical.** See also **Table S4.**

**A**

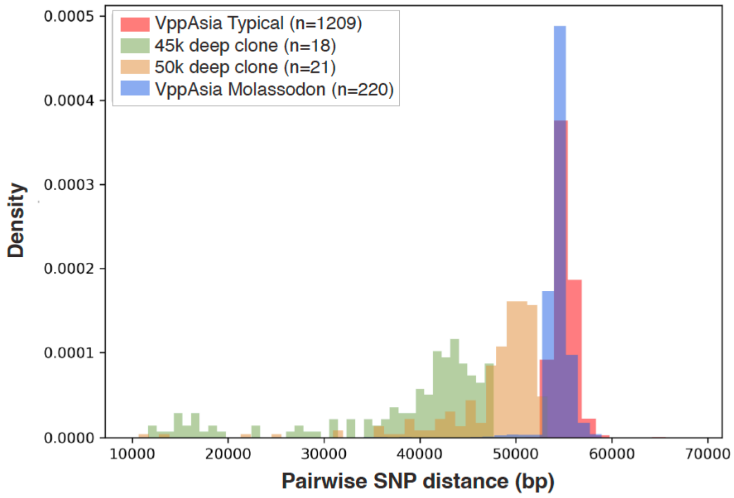


**B**

| Group | Mean SNP distance | Unrecombined genome (mean) (%) | Mean pairwise age (years) | Mean recombinations per site |
|---|---|---|---|---|
| 45 kb deep clone | 38,275 | 30.3 | 3,902 | 1.3265 |
| 50 kb deep clone | 47,713 | 13.1 | 6,571 | 2.2342 |
| Molassodon | 54,140 | 1.65 | 26,228 | 8.9175 |
| VppAsia non-redundant | 54,948 | - | - | - |

| Group | Median SNP distance | Unrecombined genome (median) (%) | Median pairwise age (years) | Median recombinations per site |
|---|---|---|---|---|
| 45 kb deep clone | 41,936 | 23.6 | 4,249 | 1.4448 |
| 50 kb deep clone | 49,512 | 9.8 | 6,840 | 2.3254 |
| Molassodon | 54,440 | 0.79 | 14,228 | 4.8375 |
| VppAsia non-redundant | 54,875 | - | - | - |

**Supplementary Figure S5. Coalescence analysis. (A)** Distribution of pairwise SNP-distances in strain subsets used for coalescence analysis. **(B)** Recombination-based estimates of divergence among deep clones and the Molassodon lineage. Recombinations per site = total number separating strains.

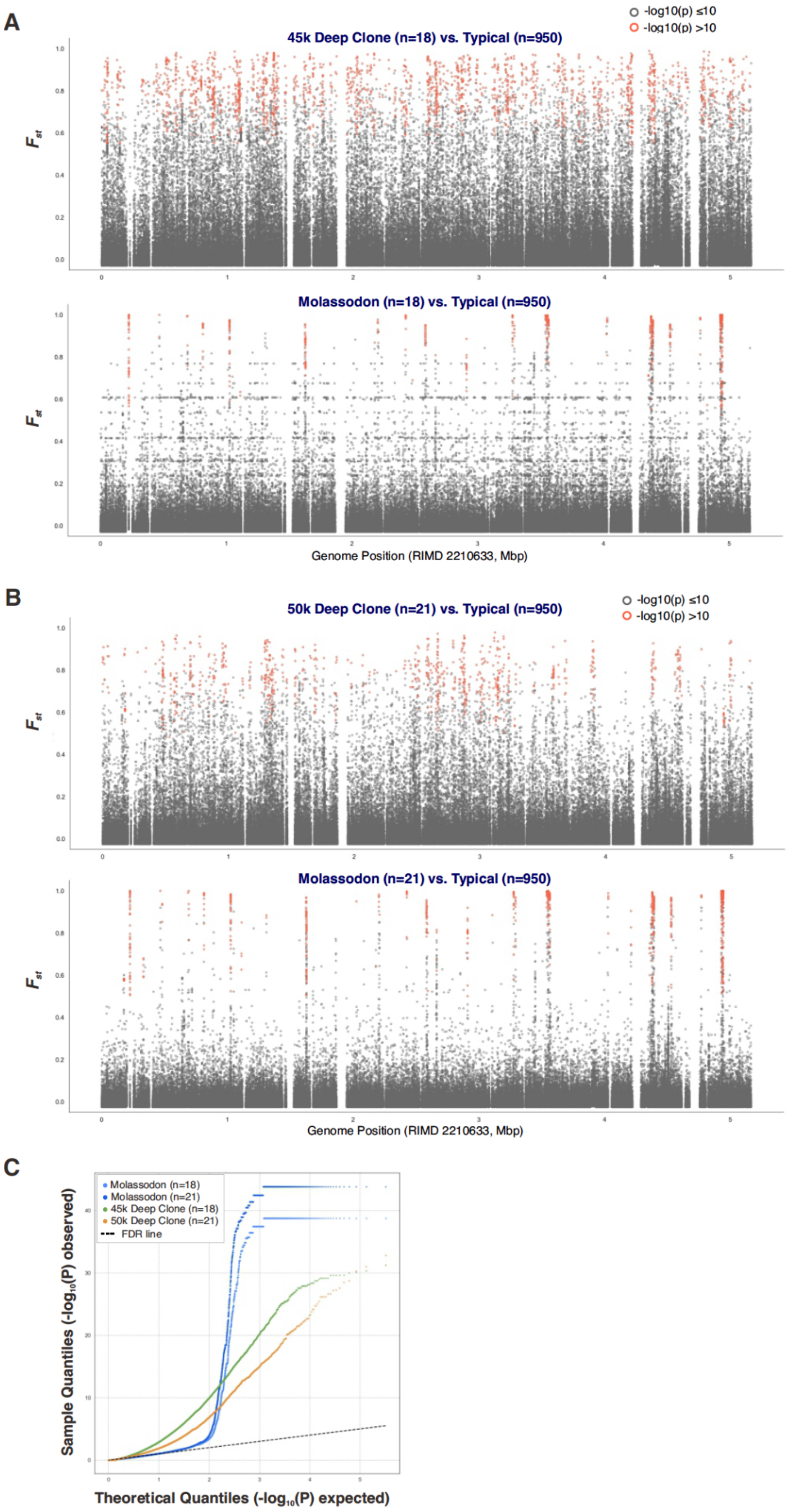


**Supplementary Figure S6. Comparison of differentiation patterns between deep clones and ecospecies.** The indicated deep clone or Molassodon datasets of the same sample size are both compared to a 950 strain VppAsia Typical strain subset. **(A)** Manhattan plots of $F_{st}$ for 18-strain deep clone and Molassodon datasets. **(B)** Manhattan plots of $F_{ST}$ for 21-strain deep clone and Molassodon datasets. For identification of the two deep clones, see **Fig. S1B** and Methods. **(C)** Q-Q (quantile–quantile) plot of $-\log_{10}(P)$ values for each comparison.

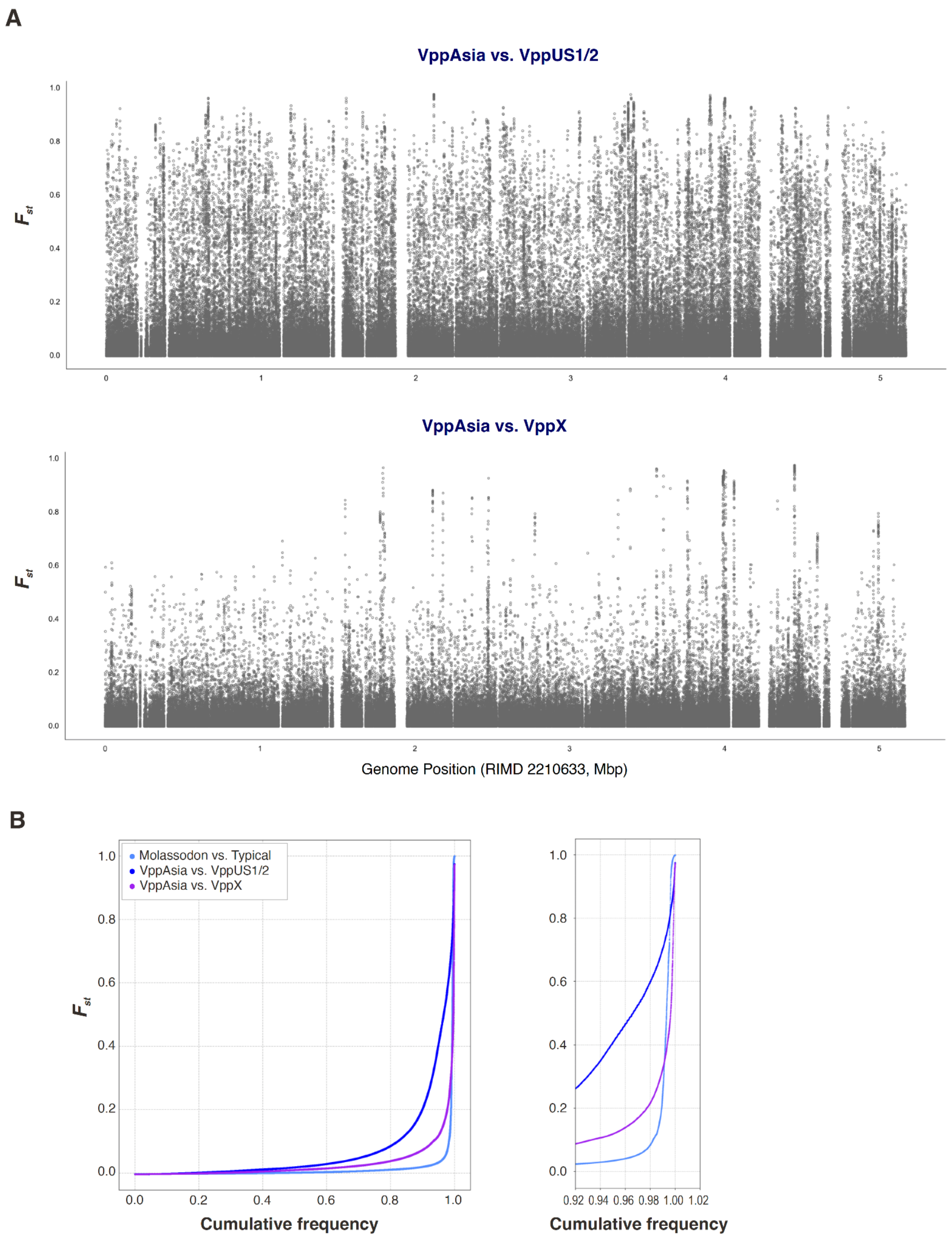


**Supplementary Figure S7. Comparison of differentiation patterns between geographic populations and ecospecies. (A)** Manhattan plots of $F_{st}$ across the reference genome calculated between the VppAsia population (n=1209) and either combined VppUS1/2 (n=165, *top*) or VppX (n=176, *bottom*) strains from the non-redundant dataset. For the Molassodon vs. Typical Manhattan plot of $F_{st}$ values, see main **Fig. 1A**. **(B)** Plot of $F_{st}$ vs. cumulative frequency for different population and ecospecies comparisons. *Right*: zoom in of 0.92-1.02 region of the x-axis.

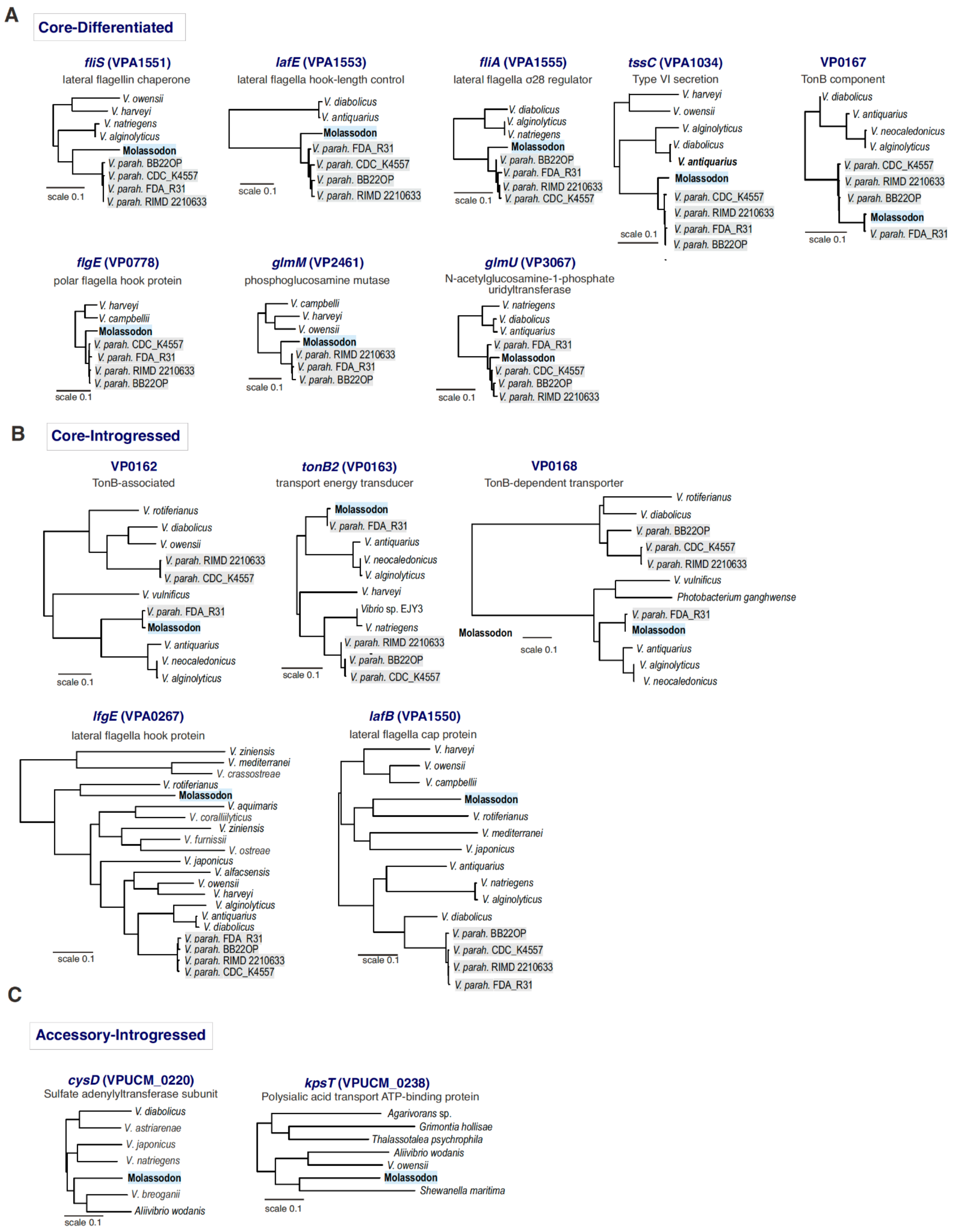


**Supplementary Figure S8. Additional trees for Molassodon accessory and differentiated trees. (A)** Example core genes that have significantly differentiated by point mutation in Molassodon (at least 5 SNPs with $F_{st}$ ≥0.5 and -log$_{10}$(p) ≥10) vs. Typical based on GWAS. **(B)** Example core genes that have significantly differentiated (at least 5 SNPs with $F_{st}$ ≥0.5 and -log$_{10}$(p) ≥10) in Molassodon showing patterns of introgression. **(C)** Example accessory genes that are fixed in Molassodon ($F_{st}$ ≥0.5 and -log$_{10}$(p) ≥10) and show patterns of introgression.

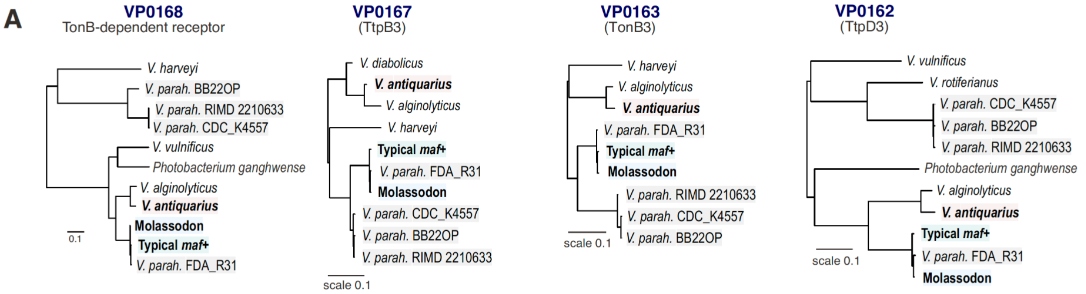


**B** VP0168

```
            1                                                                                                                              130
Molassodon  MYKNTTALSVAISLALGTAAAVAPLTAQAEEQQVEKLQKMKVTGSRISTTDMEGPSPVTVINSKDIESKGFNSAQDVLNSLTQNTGGVMAQTDSFSFTPAAQSVNLRGLGASRTLILIDGRRLPQYPLAQ
Typical maf+ MYKNTTALSVAISLALGTAAAVAPLTAQAEEQQVEKLQKMKVTGSRISTTDMEGPSPVTVINSKDIESKGFNSAQDVLNSLTQNTGGVMAQTDSFSFTPAAQSVNLRGLGASRTLILIDGRRLPQYPLAQ
Typical maf- MYKNTTALSVAISLALGTAAAVAPLTAQAEEQQVEKLQKMKVTGSRLTRASMEGSTPVAVIGRAEIERAGDVSIADVLRKSSFNSFGSHNERSGSSAQSQA-TISLRGLGSERTLVLINGKRLPGSASMG
Consensus   MYKNTTALSVAISLALGTAAAVAPLTAQAEEQQVEKLQKMKVTGSRisttdMEGpsPVtVInsk#IEskGfnSaqDVLnsltqNtgGvma#tdsfSftpaAqs!nLRGLGasRTL!LI#GrRLPqyplaq

            131                                                                                                                            260
Molassodon  NGTSNFQDIGQIPVAAIERVEVMTDSGSAIYGSDAIGGVVNFILKKDFDGVSVKVRAGDATDGDYQNGRLDVVAGKDFEDKRILFVGQFAGNEMLKQSDRDDWAGNDASKRS--QISGYSSYGANFYGDN
Typical maf+ NGTSNFQDIGQIPVAAIERVEVMTDSGSAIYGSDAIGGVVNFILKKDFDGVSVKVRAGDATDGDYQNGRLDVVAGKDFEDKRILFVGQFAGNEMLKQSDRDDWAGNDASKRS--QISGYSSYGANFYGDN
Typical maf- GGAAN---INVIPSAIVERVEVMADGGSAVYGSDAVAGVVNIILKEEFDGINVTLGSGIPSREGADEENVSIVLGTSGEKGNIMFSFEHDSKDEIYQRDRDYLSSTNTGSANYFDMSGVSIYGRNVYHDG
Consensus   nGtsNfqdIgqIPvAa!ERVEVMtDsGSA!YGSDA!gGVVNfILKk#FDG!sVkvraGdatdgdy##grld!VaGkdfEdkrI$Fvg#fagn#mlkQsDRDdwagn#askrs..#iSGySsYGaNfYgDn

            261                                                                                                                            390
Molassodon  GTH-LTPADVGSSCKEVVGEAGVDRADGK---CGYNRSADRTLKPENKQVDLLLRGEMDINDNVTAMAEARFGYKSTESQFEPNPYSVIVDDTHFRGAGEYTR--------------RMVEFGPRESNTE
Typical maf+ GTH-LTPADVGSSCKEVVGEAGVDRADGK---CGYNRSADRTLKPENKQVDLLLRGEMDINDNVTAMAEARFGYKSTESQFEPNPYSVIVDDTHFRGAGEYTR--------------RMVEFGPRESNTE
Typical maf- QLKALKGYDTDASCDPSKGFVGLTNYPGLGDICGYDYTSEAAQTASLERNTVFMNGNIFLSDDTTFNAQVLLNRNESFGRFAPAAGYFEVDPTTTGGAAFFAENGLDATKGPAAVYYRFNNVGTRDNSVT
Consensus   gth.LtpaDvgsSCkevvGeaGvdradGk...CGY#rsa#rtlkpenkqvdll$rG#mdinD#vTamA#arfgykstesqFePnpysviVDdThfrGAge%tr..............Rmv#fGpR#snte

            391                                                                                                                            520
Molassodon  ADSFGLTFGLNGLLADAYDWDV--SAGYS---EQTVETSNPAIYLTLDEAVQNGTVDLLGPISQDVVNQYSGTSTKEAYSKLYSLNGSLSGDLIELPAGMSGFAVYGEWNRTDYNESIDATTAAGGFSGL
Typical maf+ ADSFGLTFGLNGLLADAYDWDV--SAGYS---EQTVETSNPAIYLTLDEAVQNGTVDLLGPISQDVVNQYSGTSTKEAYSKLYSLNGSLSGDLIELPAGMSGFAVYGEWNRTDYNESIDATTAAGGFSGL
Typical maf- DFQADLKAGLDGTLYTDSFGEVIWETGYHLNFSNSNETGTGYVFRSPAEQLANSGQFVNGEFSADATSQLSAGTGRETQMEMHQVYGGLQFDLAEVGNIVIPLYVGAEYTTYDYFDQYDPQSEAGNIIGS
Consensus   adsfgLtfGL#GlLadaydw#V..saGYs...e#tvETsnpa!%ltldEavqNgtvdllGpiSqDvvnQySgtstkEaysk$yslnGsLsgDLiElpagmsgfaVygEwnrtDYn#siDattaAGgfsGl

            521                                                                                                                            650
Molassodon  GGTSGGGDRDQIGLGVEALIPIIEDLEMTIAGRYDHYFDDSATGGAFTPKVALAYRPIDTLLLRGSYGLGFRAPDLKRLFGEQTRGFGSGFDPVLCAQAGGSGPQDKDKYPECDTQYFDTLNGPNSELKE
Typical maf+ GGTSGGGDRDQIGLGVEALIPIIEDLEMTIAGRYDHYFDDSATGGALTPKVALAYRPIDTLLLRGSYGLGFRAPDLKRLFGEQTRGFGSGFDPVLCAQAGGSGPQDKDKYPECDTQYFDTLNGPNSELKE
Typical maf- SGNSAGGDRETYAIYAESLIAFTDELEMNIAARYDHYSD---FGDEISPKVSFRYQPLDNLMFRTGAGLGFRAPTLSDLYGADSFSSDYAKDYVFCSQNGISAS-------ECPETQYDVTRTSNEDLDA
Consensus   gGtSgGGDR#qiglgvEaLIpii##LEMtIAgRYDHYfDdsatGga.tPKValaYrPiDtL$lRgsyGLGFRAPdLkrL%Ge#trgfgsgfDpVlCaQaGgSgpqdkdkypECdtqy%DtlngpNs#Lke

            651                                                                                                                            780
Molassodon  ETSTNINLGLAWEPVDDLGLTVDWYHIKMEDVVTAPSYQDVIN-DPGKYPGTEVVRNADGTIKSVSYGPVNQSYEERSGIDLAVTYAYDTNEFGSFNTRVAVTKILIAEYQENDSEPVTDETDFMPEYTA
Typical maf+ ETSTNINLGLAWEPVDDLGLTVDWYHIKMEDVVTAPSYQDVIN-DPGKYPGTEVVRNADGTIKSVSYGPVNQSYEERSGIDLAVTYAYDTNEFGSFNTRVAVTKILIAEYQENDSEPVTDETDFMPEYTA
Typical maf- EKSVSFNFGISYSPIDDLDLTLDYYNISIDDVITLNTLQSMIDQERSSGVSNPNIVRENGRIKEATAGLQNLGTLDTSGLDLKIGYRYDFN-VATIRYDFTGSYVLDYSSPEYVGGPVNNQVGRNGLPEY
Consensus   EtStniNlGlaweP!DDLgLTvDwYhIkm#DV!TapsyQdvI#.#pgkypgtev!rna#GtIKsvsyGpvNqsye#rSGiDLa!tYaYDtNefgsfntrvavtk!LiaeyqEndsePVt##tdfmpeyta

            781                                                                                                               892
Molassodon  TLNLGWAYEDVAVNLFGKYRDRMCSSYANRYYFDDCDAAKKAGYKAEIASMTTWNLTASYQFNESGRVTVGAINMFDKTPPADPLSDTSPFYANGYDDPIGRQLYIEASYDF
Typical maf+ TLNLGWAYEDVAVNLFGKYRDRMCSSYANRYYFDDCDAAKKAGYKAEIASMTTWNLTASYQFNESGRVTVGAINMFDKTPPADPLSDTSPFYANGYDDPIGRQLYIEASYDF
Typical maf- RFNTG-----VGVNFLEEHDIYLSADHIADQAQDVDDNYNKTGH---ISSQTTWNIAYNYLAPWDAKFTAGVRNLTDEDPAFE--SDGVTYDKDLYSIQ-GRVYFLKYSQNF
Consensus   tlNlGwayedVaVNlfgkyrdr$cssyanryyfDdcDaakKaGykaeIaSmTTWNltasYqfnesgrvTvGaiN$fDktPpa#plSDtsp%ya#gYddpiGRql%ieaSy#F
```

**Supplementary Figure S9. Differentiation at the VP0162-VP0168 *tonB3* locus in Molassodon and *maf*+ Typical strains. (A)** Phylogenetic relationship between differentiated alleles at the *tonB3* locus for Molassodon, Typical *maf*+/- strains, and KEGG homologs from other species. *V. parahaemolyticus* strain FDA_R31 is a rare Typical *maf*- strain that carries the Molassodon/-like *tonB*-locus alleles. **(B)** Alignment of protein sequences for VP0168 homologs, encoding TonB-dependent receptors, generated with multalin [1].

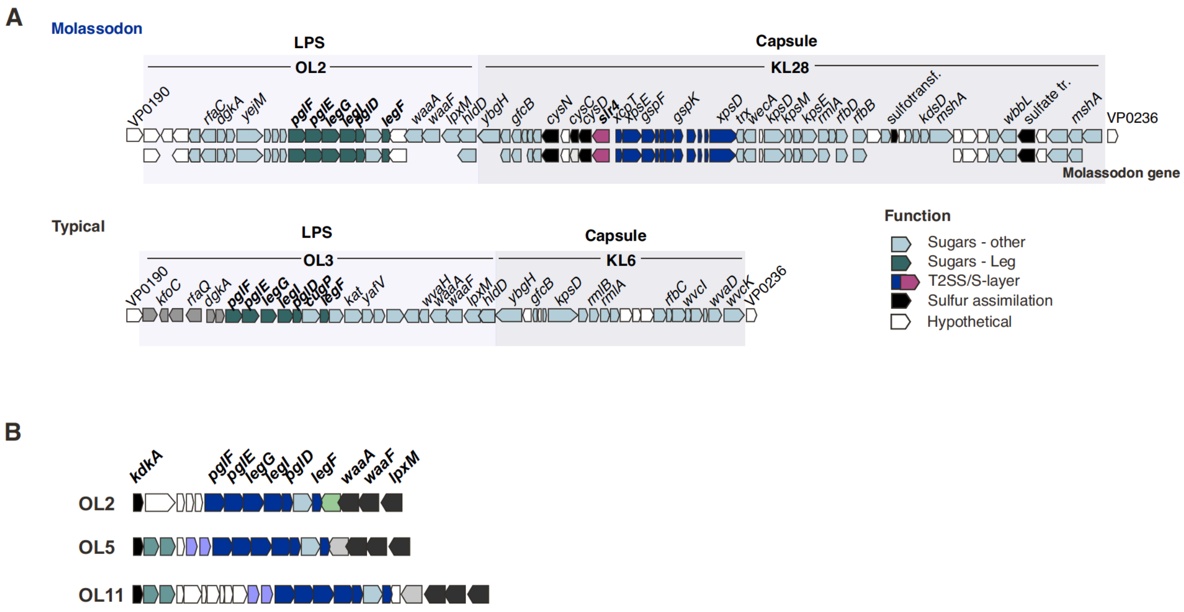


**Supplementary Figure S10. Comparison of LPS/capsule islands and serotypes between Molassodon and Typical strains. (A)** Schematic maps of example LPS/capsule loci for Molassodon and Typical strains. For Molassodon, the most commonly detected serotype in our dataset is shown (OL2:KL28). For Typical, the OL3:KL6 locus of the reference strain RIMD 2210633 is shown. **(B)** Synteny of O-serotype genes identified in Molassodon strains (139 strain dataset). Based on Kaptive annotation [2]. Similar genes are in the same colour, while unrelated genes are white. Putative Leg (legionaminic acid) biosynthesis genes are in dark blue.

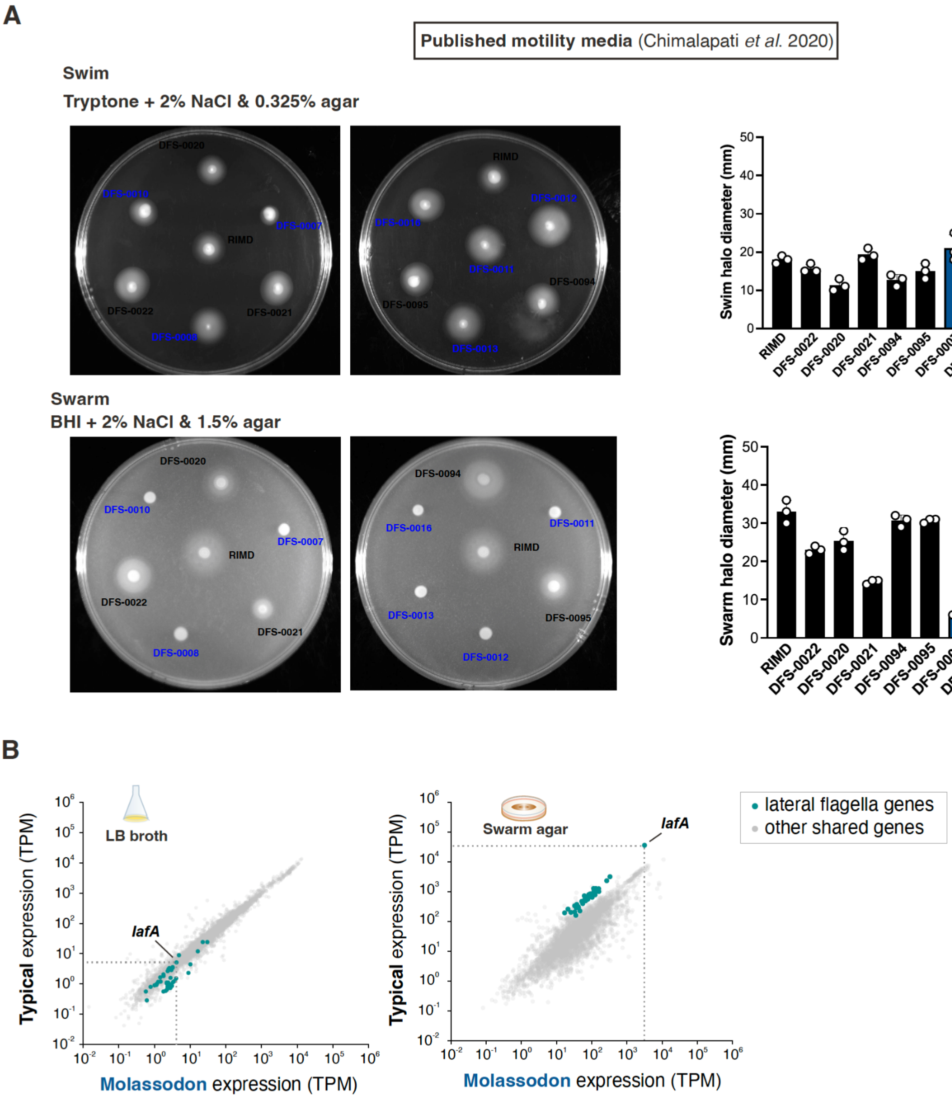


**Supplementary Figure S11. Motility phenotypes and expression of natural Typical and Molassodon strains with published *V. parahaemolyticus* media. (A)** Swimming and swarming tested using published media compositions [3]). Assays were performed in parallel for 12 hours at room temperature. **(B)** RNA-seq expression in Molassodon vs. Typical bacteria in broth culture (LB, *left*) and on swarm agar (*right*). Typical cells were recovered from the outside of swarm halos, while Molassodon RNA was isolated from bacteria on the edge of the macrocolony that forms on swarm agar. Green dots: all lateral flagella genes (differentiated and undifferentiated). Expression is plotted in TPM (transcripts per million).

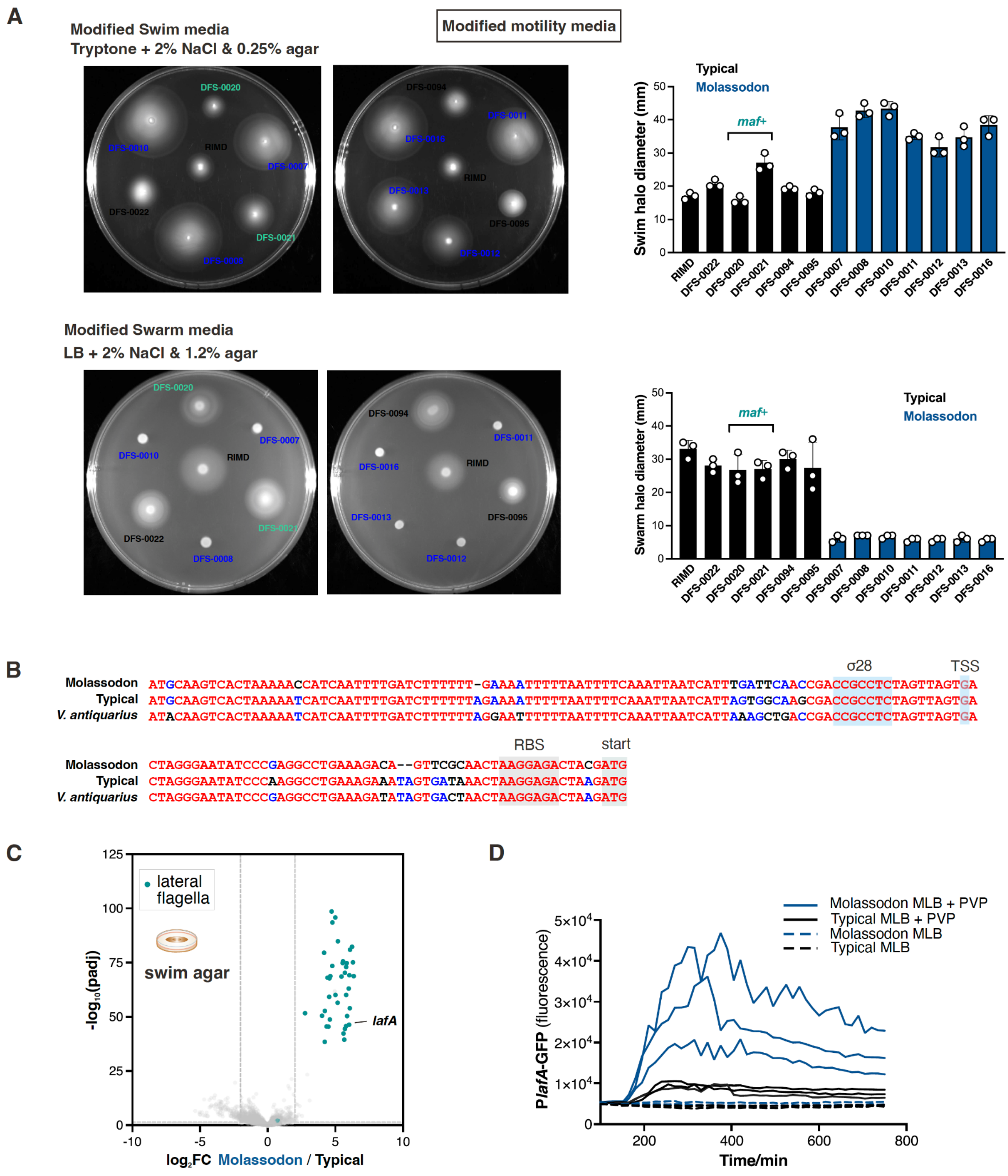


**Supplementary Figure S12. Motility phenotypes and expression of natural Typical and Molassodon strains in modified assays. (A)** Swimming and swarming tested in modified media compositions. Assays were performed in parallel for 12 hours at room temperature. **(B)** Alignment of the regions upstream of the *lafA* start codon from a Molassodon strain, Typical strain, and *V. antiquarius* EX25. Red: 100% conserved. Blue: conserved in 2/3 strains. Black: not conserved. RBS: putative ribosome binding site. σ28: putative sigma-28 (LafS) binding site. start: start codon. Generated with multalin [1]. **(C)** RNA-seq differential expression of shared genes in the swimming condition for three Molassodon vs. Typical strains. Green: all lateral flagella genes (differentiated and undifferentiated). **(D)** Expression of respective P*lafA* reporters in three Molassodon and Typical strains in the absence (MLB) and presence (MLB+3% PVP) of increased viscosity provided by polyvinylpyrrolidone (PVP). Each line represents the mean of two technical replicates for three Typical (black lines) or three Molassodon strains (red lines) in MLB (dashed lines) or MLB + 3% PVP (solid lines). PVP: polyvinylpyrrolidone. Related to main **Fig. 2F.**

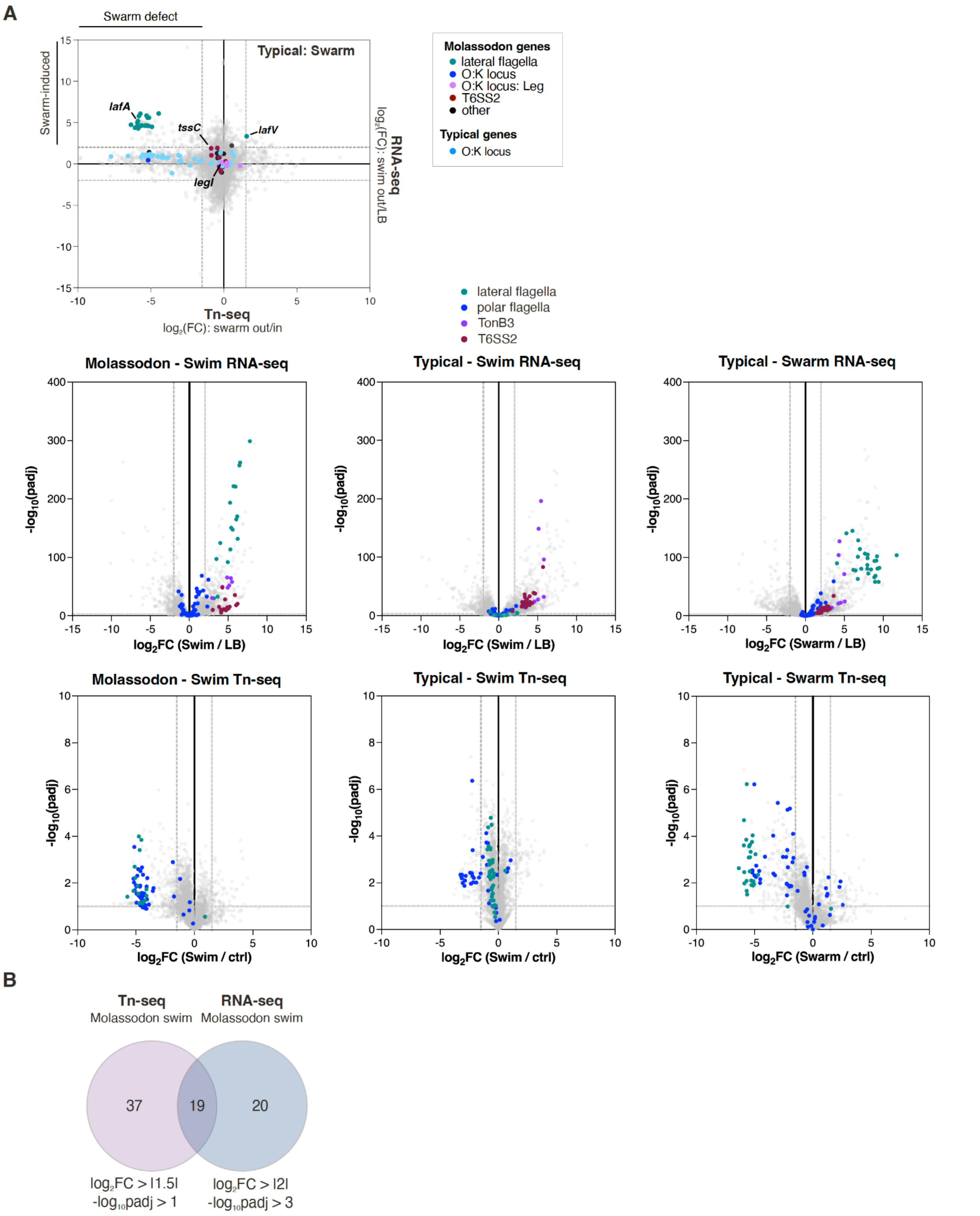


**Supplementary Figure S13. Additional data related to Tn-seq/RNA-seq. (A)** *Top:* Tn-seq and RNA-seq profile for the Typical strain during swarming. Tn-seq: RIMD 2210633, $log_2FC$ Swarm halo vs. Swarm inside. RNA-seq: mean of three Typical strains, $log_2FC$ Swarm halo vs. LB. Molassodon genes identified by GWAS are coloured. Labeled Typical genes (light blue) are encoded in the OL3 LPS locus. All other genes are grey. *Bottom:* Volcano plots for RNA-seq (top, data for three strains) and Tn-seq (bottom, data for one strain). Cutoffs are as follows: RNA-seq, $log_2FC > |2|$ and $-log_{10}padj > 3$ / Tn-seq, $log_2FC > |1.5|$ and $-log_{10}padj > 1$. n=3. **(B)** Overlap of significantly induced / required genes for Molassodon swimming.

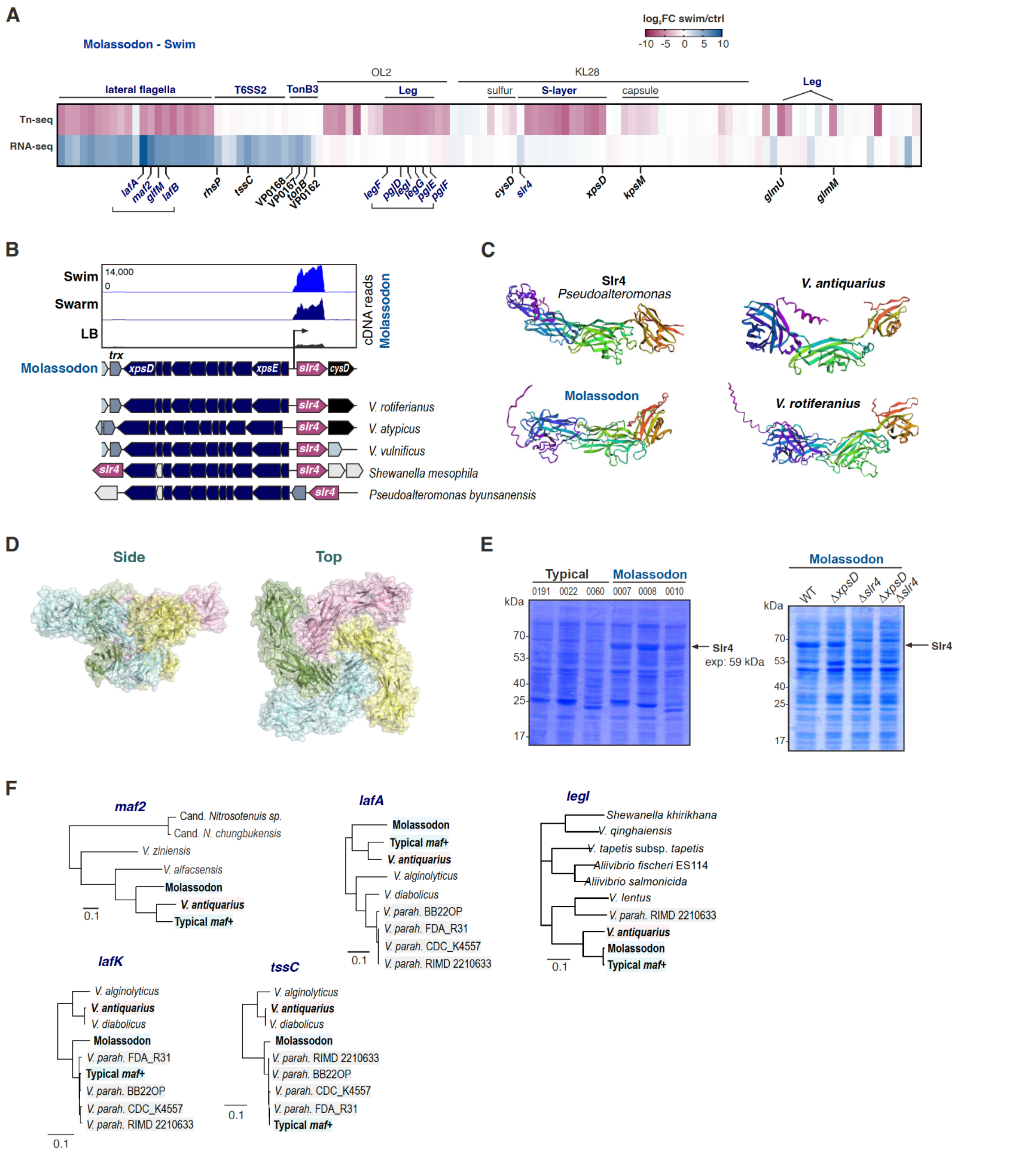


**Supplementary Figure S14. Swim phenotype and expression links additional traits to the Molassodon phenotype. (A)** Tn-seq phenotype and RNA-seq expression for all Molassodon genes in Molassodon strain DFS-0010 in the swim condition. **(B)** RNA-seq expression (*top*) and synteny analysis (*bottom*) for the putative Slr4 S-layer protein homolog and adjacent Type II secretion system (T2SS) in Molassodon and other species. Bent arrow: putative transcription start site. Blue genes: putative T2SS components. Dark pink: Slr4. cDNA coverage was visualized with Integrated Genome Browser (IGB) [4]. Synteny analysis was performed with webflags [5]. **(C)** Predicted tertiary structures of Slr4 from *Pseudoalteromonas* and Molassodon. **(D)** Predicted tetrameric complex formed by Molassodon Slr4. Structures were predicted with Alphafold2/Alphafold Multimer at Colabfold, [6,7]) and visualized with Pymol. **(E)** 1D SDS-PAGE analysis of total proteomes from Typical WT, Molassodon WT, and Molassodon deletion mutant strains. *xpsD*: gene encoding Type II secretion component (see panel A). *slr4*: gene encoding putative S-layer protein. **(F)** Relationship between KEGG orthologs from other species for Molassodon, Typical *maf*-, and Typical *maf*+ versions for differentiated accessory (*maf2*, *legI*) and core (*tssC*, *lafK*, *lafA*) genes.

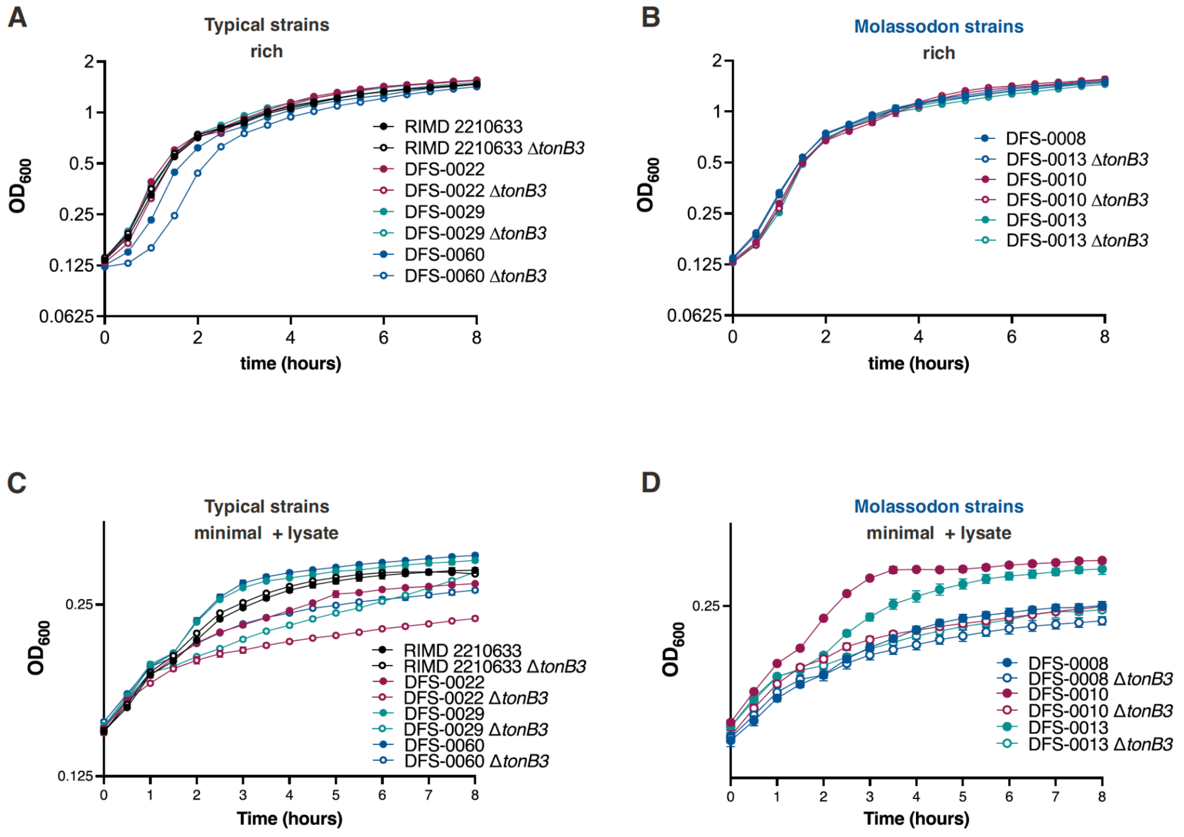


**Supplementary Figure S15. Minimal medium + lysate growth curve data for individual strains. (A)** Typical strains. **(B)** Molassodon strains. n=3. Related to main **Fig. 3E.**

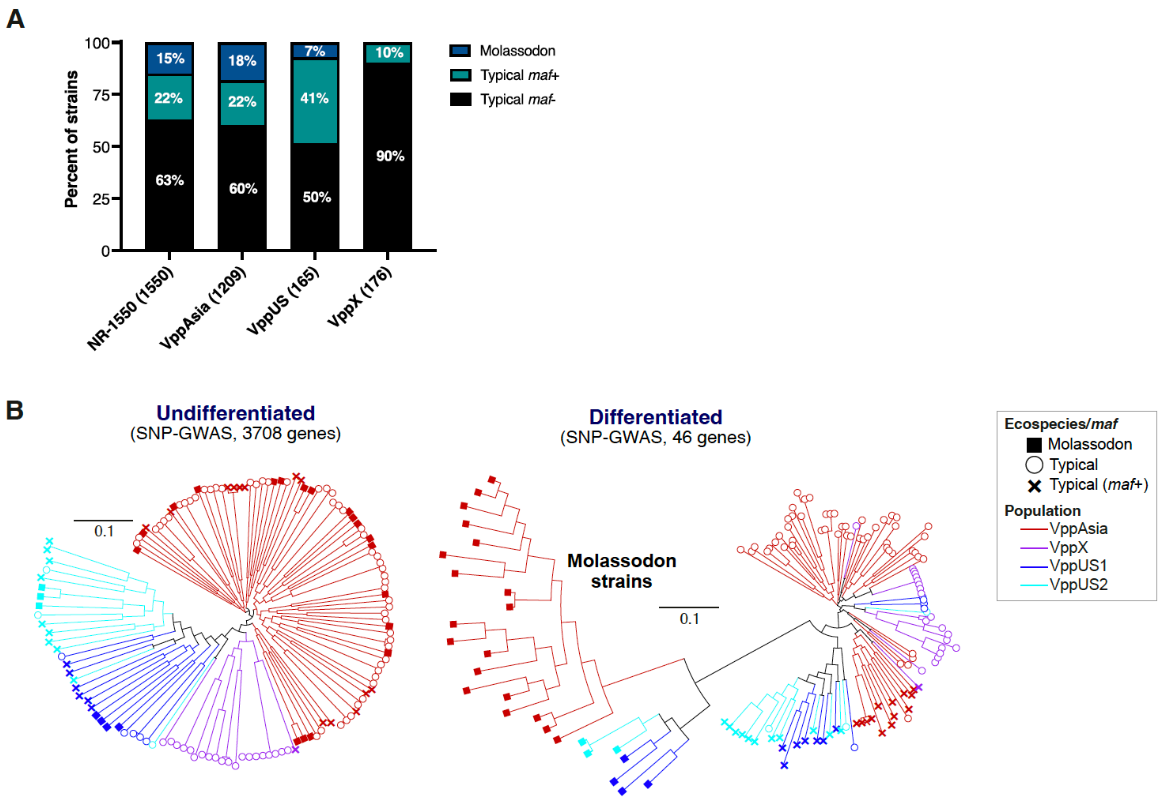


**Supplementary Figure S16. Distribution and phylogenetic relationships between Molassodon, Typical *maf*+/*maf*- for Undifferentiated and Differentiated Molassodon genes. (A)** Distribution of Molassodon and Typical *maf*-/*maf*+ strains in the non-redundant dataset used to measure differentiation. **(B)** Approximate Maximum Likelihood trees for Undifferentiated (top, 3708 genes) and Differentiated genes (bottom, 46 genes) identified in core genome SNP GWAS. A subset of 137 strains with representatives of each group of interest (population, ecospecies) was used. Differentiated genes: at least five differentiated SNPs. Undifferentiated genes: no SNPs with Fst ≥0.05. Note that two highly differentiated core genes identified in the pan-GWAS are not included. Branches: geographic population. Tips: ecospecies (Typical/Molassodon) and *maf* (*maf2-glfM* accessory genes) genotype.

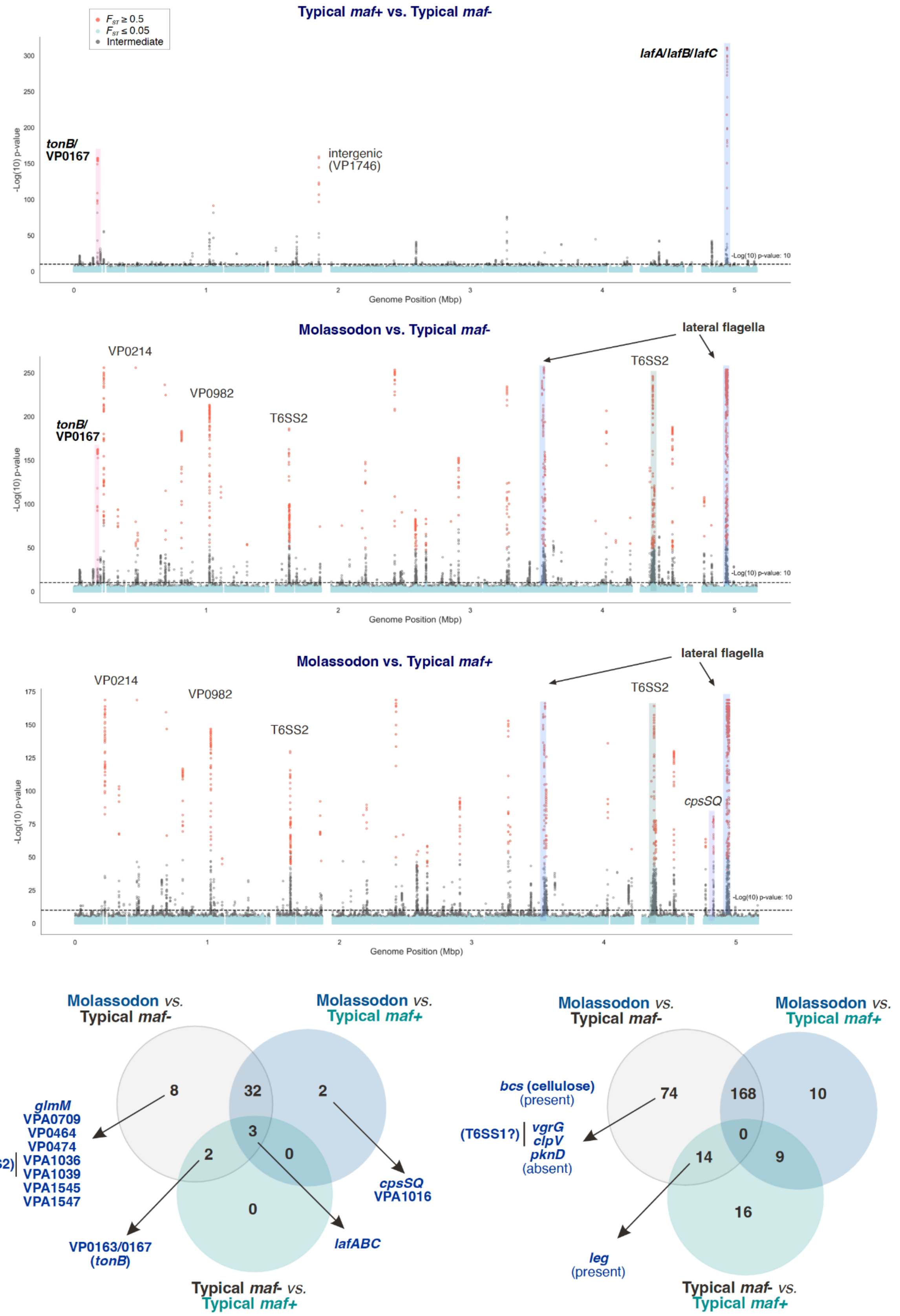


**Supplementary Figure S17. Typical *maf+* differentiation.** See also **Table S2.** *Top*: Manhattan plots for differentiation between Molassodon, Typical *maf*+, and Typical *maf*-. *Bottom*: Overlap of core and accessory differentiated genes between the three GWAS in panel A. Differentiated = -log10p >10 and $F_{ST}$ > 0.5.

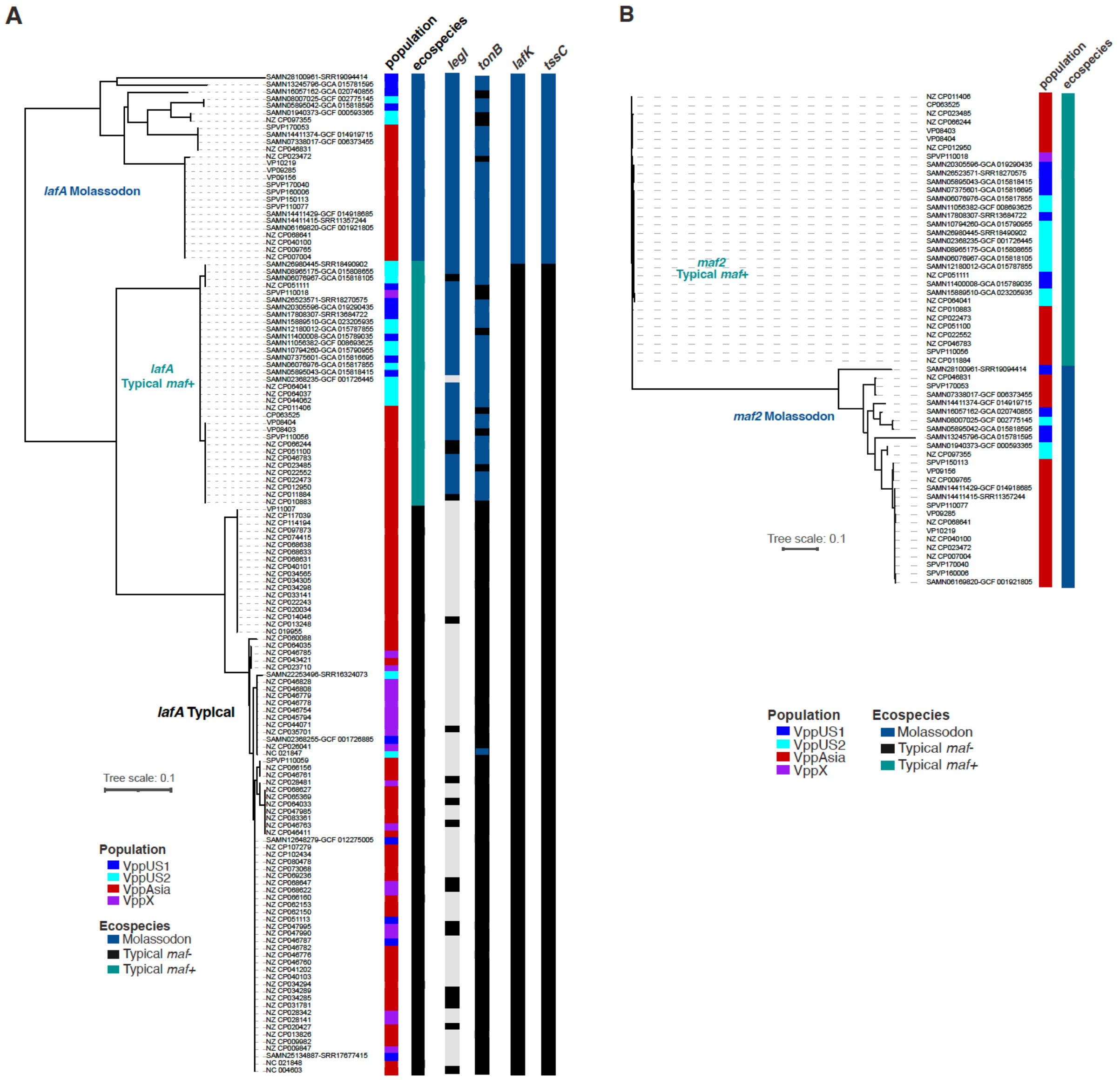


**Supplementary Figure S18. Phylogenetic relationship between Molassodon core and differentiated genes in *V. parahaemolyticus*. (A)** Neighbour-joining tree for lateral flagella gene *lafA* (VPA1548). Allelic variants of *maf2*, *legI*, *tonB*, *lafK*, and *tssC* based on phylogenetic trees (Fig. 18B; Fig. 19) are indicated. **(B)** Neighbour-joining tree for lateral flagella accessory gene *maf2*.

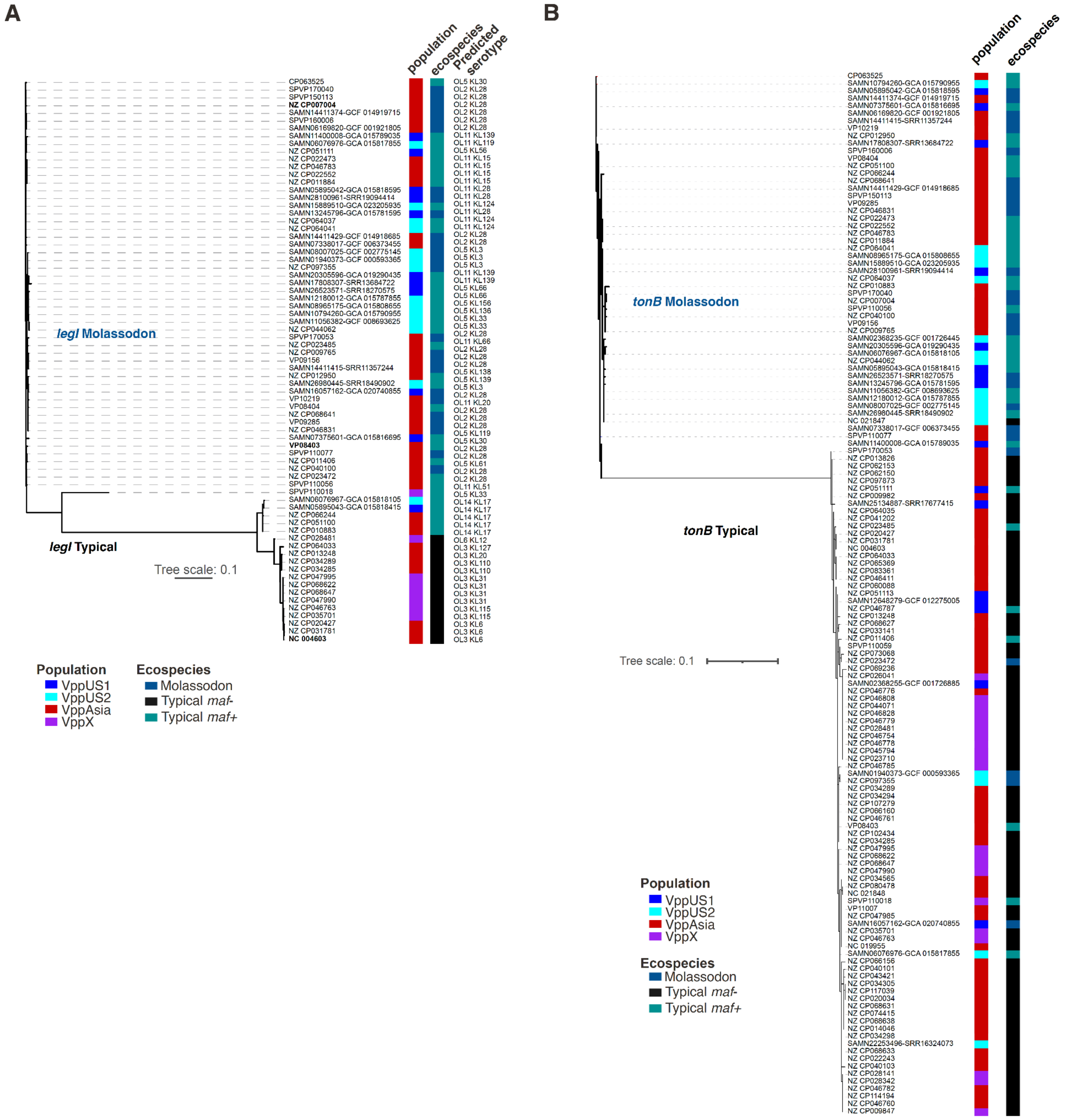


**Supplementary Figure S19. Phylogenetic relationship between Molassodon core and differentiated genes in *V. parahaemolyticus*. (A)** Neighbour-joining tree for accessory legionaminic acid biosynthesis gene *legI*. **(B)** Neighbour-joining tree for transport gene *tonB* (VP0163).

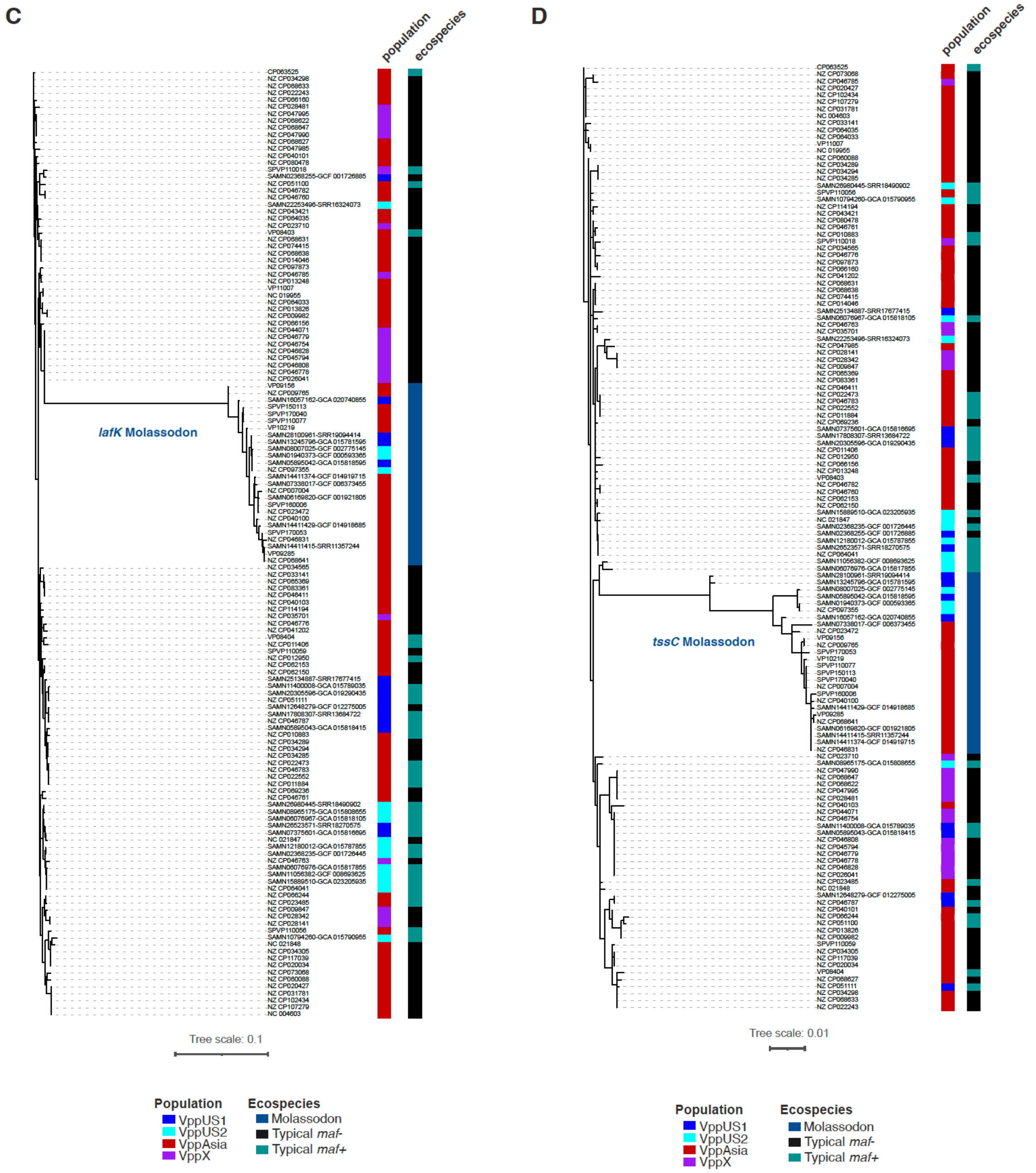


**Supplementary Figure S19 (continued). Phylogenetic relationship between Molassodon core and differentiated genes in *V. parahaemolyticus*. (C)** Neighbour-joining tree for lateral flagella core gene *lafK* (VPA1538). **(D)** Neighbour-joining tree for Type VI secretion system 2 gene *tssC* (VPA1034).

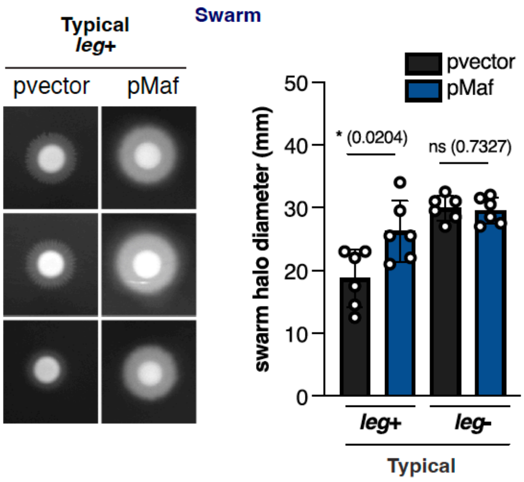


**Supplementary Figure S20. Swarming after heterologous expression of *maf* from Molassodon into Typical *maf*- strains.** * $p$<0.05, ** $p$<0.01, ns: not significant, Student's $t$-test.

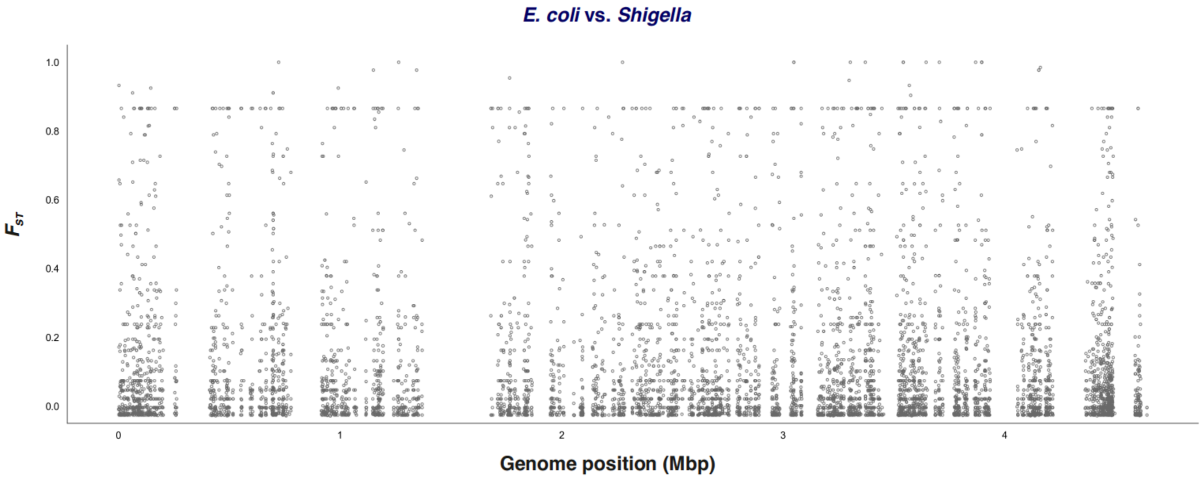


**Supplementary Figure S21. Core genome $F_{ST}$ analysis between *E. coli* and *Shigella* strains.** A previously published non-redundant dataset was used [8].

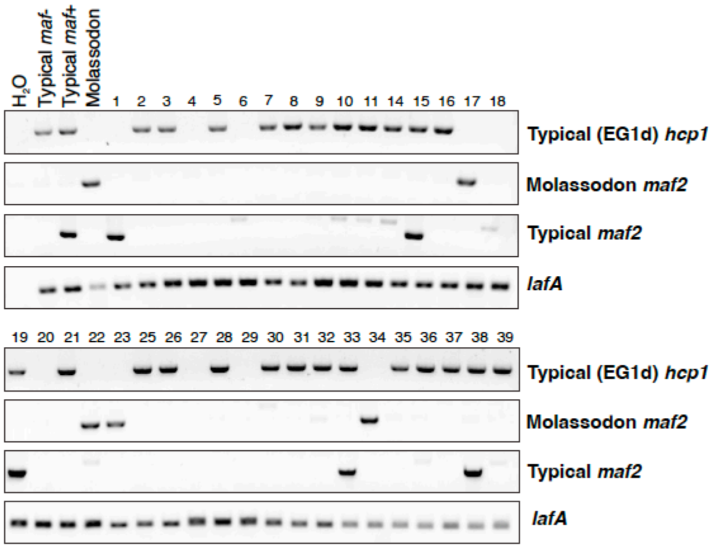


**Supplementary Figure S22. Detection of *V. parahaemolyticus* Molassodon and Typical (*maf*- and *maf*+) strains in a single sample of seawater.** Colonies were recovered on ChromAgar from seawater samples from Fujian Province, China (平潭 Pingtan and 泉州 Quanzhou). Mauve colonies on CHROMagar™ *Vibrio* plates were confirmed to be *V. parahaemolyticus* by PCR using primers binding *lafA* in both Molassodon and Typical strains (DFO-0152/0153). Molassodon strains were identified with primers binding the Molassodon version *maf2* (DFO-0148/0149) and intermediate strains with primers binding the Typical version of *maf2* (DFO-1003/1004), while EG1d (see was verified with primers binding *hcp1* (VP1393, DFO-0378/0379).
